# Maladaptation in cereal crop landraces following a soot-producing climate catastrophe

**DOI:** 10.1101/2024.05.18.594591

**Authors:** Chloee M. McLaughlin, Yuning Shi, Vishnu Viswanathan, Ruairidh Sawers, Armen R. Kemanian, Jesse R. Lasky

**Author notes:** **Classification**: Biological Sciences (Evolutionary Biology).

## Abstract

Aerosol-producing global catastrophes such as nuclear war, super-volcano eruption, or asteroid strike, although rare, pose a serious threat to human survival. Light-absorbing aerosols would sharply reduce temperature and solar radiation reaching the earth’s surface, decreasing crop productivity including for locally adapted traditional crop varieties, i.e. landraces. Here, we test post-catastrophic climate impacts on barley, maize, rice, and sorghum, four crops with extensive landrace cultivation, under a range of nuclear war scenarios that differ in the amount of black carbon aerosol (soot) injected into the climate model. We used a crop growth model to estimate gradients of environmental stressors that drive local adaptation. We then fit genotype environment associations using high density genomic markers with gradient forest offset (GF offset) methods and predicted maladaptation through time. As a validation, we found that our GF models successfully predicted local adaptation of maize landraces in multiple common gardens across Mexico. We found strong concordance between GF offset and disruptions in climate, and landraces of all tested crop species were predicted to be the most maladapted across space and time where soot-induced climate change was the greatest. We further used our GF models to identify landrace varieties best matched to specific post-catastrophic conditions, indicating potential substitutions for agricultural resilience. We found the best landrace genotype was often far away or in another nation, though countries with more climatic diversity had better within-country substitutions. Our results highlight that a soot-producing catastrophe would result in the global maladaptation of landraces and suggest that current landrace adaptive diversity is insufficient for agricultural resilience in the case of the soot scenarios with the greatest change to climate.

## Introduction

Environmental variability due to changing climate poses one of the greatest threats to agricultural productivity^1^. Increasingly, researchers aim to predict the effects of changing climate on agriculture, projecting constraints on crop production and anticipated decreases in yield^2,3^. For regions and crop species identified as vulnerable under future climates, strategies to increase agricultural resilience may include adapting management practices and substituting varieties or crop species^4^.

A catastrophic incident is defined by the National Response Framework as, “any natural or manmade incident, including terrorism, that results in extraordinary levels of mass casualties, damage, or disruption severely affecting the population, infrastructure, environment, economy, national morale, and/or government functions”. Aerosol-producing global catastrophic events, such as nuclear war, asteroid strike, or super-volcano explosion, are expected to produce significant climate change^5^ through deflecting solar radiation, preventing sunlight from reaching the Earth’s surface and causing global cooling. Since the spread of nuclear weapons during the twentieth century, there has been significant focus on assessing the consequences of a nuclear conflict on both society and the environment^6^. Published climate models have been used to consider the impacts of nuclear wars on the growth of major grain crops^7–9^ and summarize the degree to which the rapid environmental change induced by a black carbon aerosol (soot) producing catastrophe would impact global crop production. To date, the impact of such a soot-producing catastrophe on agricultural systems has not accounted for intraspecific diversity present in crop species, including landraces, and how this diversity may aid in increasing agricultural resilience. Cereal crops account for the most calories consumed by humans^10^ and maintaining their production post-global catastrophe is of the utmost importance.

Crop landraces (local traditional varieties) contain most of the genetic diversity within many crops, much of which is not represented in modern breeding varieties^11^ and are still widely cultivated in the developing world. The continual cultivation and selection of crops by farmers gives rise to these local varieties that often carry locally adapted alleles and phenotypes^12^. Historically, landraces have contributed to plant breeding through the identification of traits and alleles for adaptation to stressful environments (e.g. water stress, salinity, and high temperatures)^13^. Many thousands of landrace varieties are now stored in germplasm banks and represent untapped adaptive diversity that may increase agricultural resilience under changing environments^14^.

The genetic basis of adaptation to local environments can be characterized through geographic associations between genotype and environment, known as genotype-environment associations^15^. Genotype-environment associations have been used to study the adaptive potential of species^16^, estimate optimal range shifts^17^, and identify genes that may be advantageous for organisms under future climates^18^. Genotype-environment associations may also give insights into which specific environmental pressures drive local adaptation^19–21^. For landraces, a large portion of genomic variation can be explained by environments of origin^22–24^, making them good systems for considering the environmental gradients driving local adaptation^25^ and the geographic distribution of locally adapted alleles^26,27^.

An emerging approach for predicting adaptation to novel environments is first fitting genotype-environment models that describe how current allele frequencies change across environments under an assumption of local adaptation. Next, the fitted model is applied to a novel environment to determine the change in genomic composition required for adaptation to that environment, known as genomic offset (reviewed in ^28^). The genotype-environment models can further be extended to identify optimal genotypes or varieties for specific environments^24,29^ and guide movement of genotypes to minimize maladaptation to the novel climates. Such modeling methods capture long-term signals of adaptation and may provide insights into genotypes that are the most vulnerable/sources of resilience to climatic variability^30^.

We studied the climate impacts of a soot-producing catastrophe on broadly distributed globally important cereal crops for which landrace cultivation is important for smallholder farmers: *Sorghum bicolor* (L.) Moench (sorghum), *Zea mays* L. (maize), *Oryza sativa* L. subsp. *indica* and *japonica* (rice), and *Hordeum vulgare* L. (barley). These crops represent four of the top five cereal species in global production^31^. For each crop species included in this study, we independently implemented crop growth models to identify environmental stressors and genomic models to estimate the degree of disruption to current landrace adaptation under several post-catastrophic scenarios differing in the amount of soot injected into the climate model. We validated our genomic models through comparing predicted local adaptation and published maize landrace performance data collected in common gardens across diverse climates in Mexico. We further extended our genomic models to identify landrace varieties best matched to specific post-catastrophic conditions, supporting the management strategy of substituting vulnerable landrace genotypes for more resilient ones.

Our study aims to evaluate the environmental forces that have historically shaped genomic variation in landraces and to assess how landrace adaptation may be disrupted by novel catastrophic events. There is little research investigating the impacts of changing climate on diverse genotypes of multiple species. Thus, the literature may be oversimplifying climate change effects on agricultural and ecological systems. Utilizing a multi-species genomics approach allows us to confront this challenge, acknowledging the distinct impacts on various species that are vital for food production. We hypothesized that the magnitude of maladaptation would be largely determined by the magnitude of environmental change, that substitutions of genotypes from different locations could partially ameliorate effects of climate change, and that countries with greater climatic diversity would have better adapted genotype substitutions within their borders compared to less climatically diverse countries. Finally, the approach developed in this study may be extended to and prove valuable for understanding impacts of greenhouse gas induced climate change.

## Results

### Climate scenarios

We studied disruptions to current landrace adaptation for six nuclear war scenarios that simulate the impact of varying amounts of stratospheric soot on global climate (Fig. 1b) using previously published climate simulation data^6,32^. The published weather files describe the climate impacts for five India-Pakistan nuclear war scenarios (soot injections of 5 Tg, 16 Tg, 27.3 Tg, 37 Tg, and 46.8 Tg), one United States-Russia scenario with a soot injection of 150 Tg, and a control run that describes normal fluctuations in climate.

**Figure 1.**
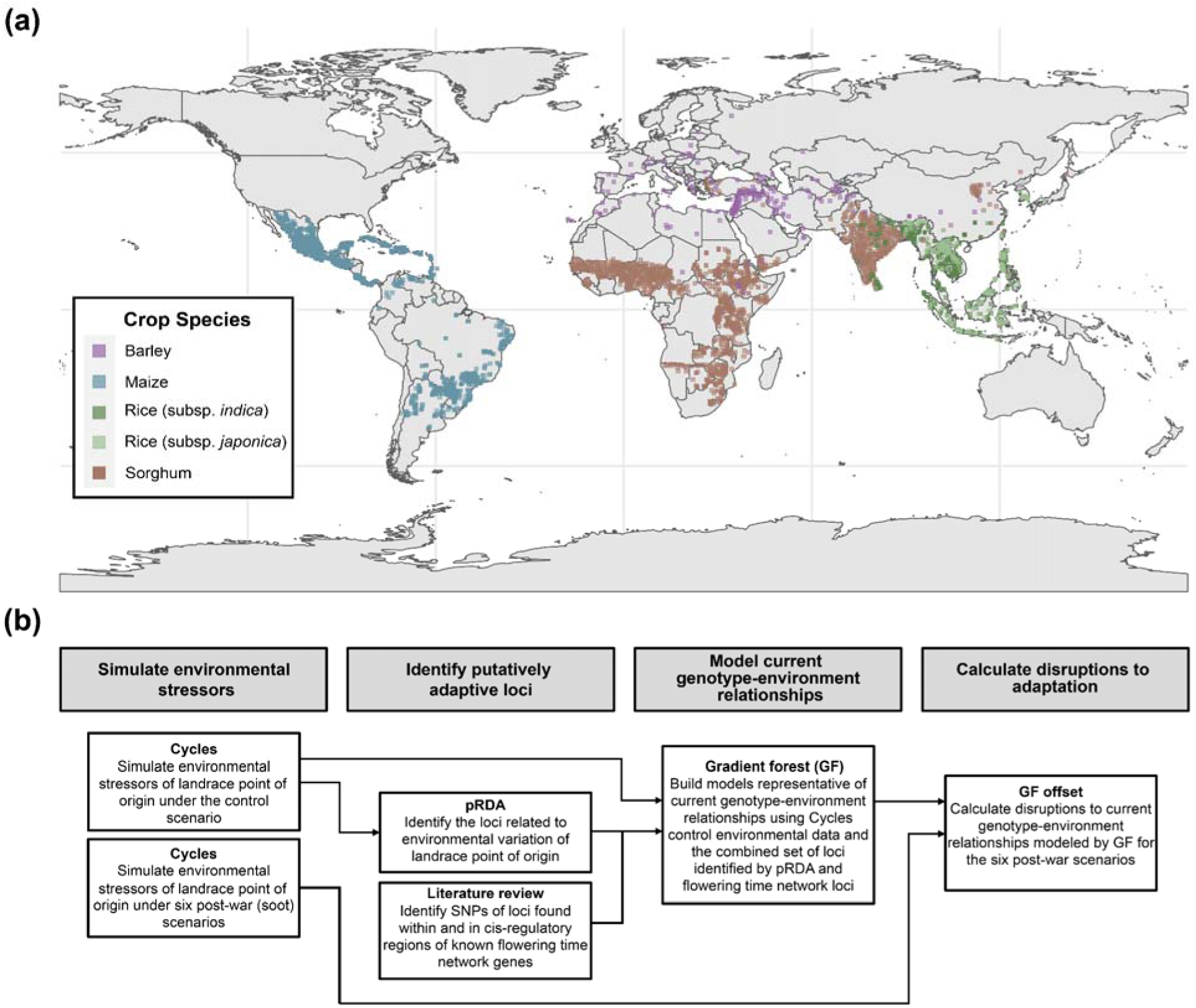
a) Global distribution of genotyped landrace accessions used in the study (All accessions, *n* = 6,384; Barley, *n* = 215; Maize, *n* = 3,404; Rice subsp. *indica*, *n* = 677; Rice subsp. *japonica n* = 309; Sorghum, *n* = 1779). b) Modeling and statistical pipeline used in this study.

### Genotyped landrace accessions

To assess maladaptation in cereal crop landraces following a soot-producing catastrophe, we identified species for which landrace relatives are currently grown in the developing world that also had publicly available, high quality sequencing data of geographically diverse accessions. From these criteria, we selected four crop species: barley (*n* = 215), maize (*n* = 3,404), rice (*n* = 677 of the subsp. *indica; n* = 309 of the subsp. *japonica*), and sorghum (*n* = 1,779). The distribution of accessions covered most of the agricultural areas in the developing world (Fig. S1a) across diverse climate regimes.

### Using crop growth models to estimate integrated local climatic stressors under control and post-war conditions

Traditional implementations of genotype-environment associations typically use off-the-shelf climate parameters without connection to organismal biology and without consideration of phenology. However, actual climate-driven stress likely emerges from a combination of conditions (e.g. precipitation and temperature) and depends on organismal phenology and development. To address these issues, we used the *Cycles* agroecosystem model^2,33^ to simulate growth and stress parameters for our full set of genotyped, georeferenced landrace accessions (*n* = 6,384) under control and six nuclear war conditions that differed in the quantity of stratospheric soot simulated (5 Tg, 16 Tg, 27 Tg, 37 Tg, 47 Tg, 150 Tg). *Cycles* simulations selected a planting date in a designated planting window when the weather and soil conditions are suitable for the specific crop, and simulated crop growth until the time of harvest or termination, using parameters specific to each of our four species. *Cycles* simulations that accounted for species-specific growth parameters were run independently for each crop species and climate scenario (control and six soot scenarios).

We used outputs from *Cycles* simulations to infer emergent environmental, growth, and stress values experienced during key phenological stages of crop, constrained to the growing period for each simulated accession under the different climate scenarios (Table S1). Thus, the selected model outputs characterized differences in environment and potential stress experienced by a given landrace accession under control and post-war climates, while accounting for crop-specific growth parameters. For each climate scenario and accession, we extracted 13 *Cycles*-derived variables representative of average temperature, coldest temperature, water stress, and solar radiation experienced by simulated landrace accessions across the vegetative, reproductive, and total growth and days to reach maturity (hereafter, *Cycles*-derived environmental variables, Fig. 2, Table S1). While in reality landrace accessions likely exhibit variation in response to environmental variability, modeling this genetic variation was not our goal at this stage. Rather, our goal was to use *Cycles* to estimate integrative environmental stressors through space and time for later use in modeling genotype-environment associations.

**Figure 2.**
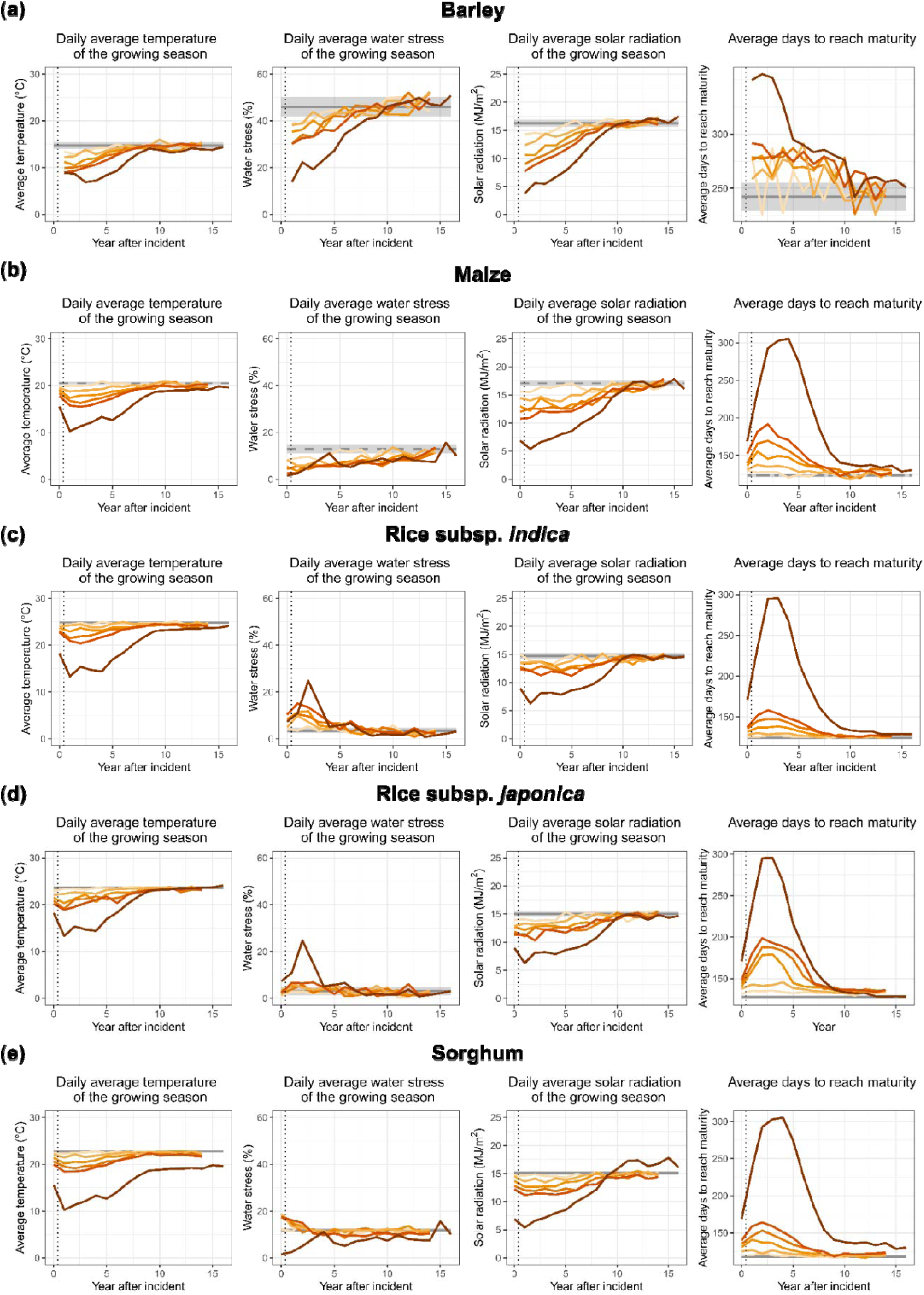
*Cycles*-derived environmental variables for A) barley B) maize C) rice subsp. *indica* D) rice subsp*. japonica* and E) sorghum. Control lines are plotted as the average value across all accessions that were projected to reach maturity. The shading around averaged control represents the standard deviation of yearly averages, indicating yearly fluctuations in environmental variables. For all soot scenarios, lines are plotted as yearly averaged values across accessions that were projected to reach maturity. The vertical dotted line indicates the time of soot injection into the climate models.

As described by other groups, in all war scenarios regardless of detonation location, produced soot spreads globally and causes disruptions to solar radiation reaching the earth’s surface, resulting in global cooling^6,32^. Stratospheric soot from each post-war scenario dissipated over the course of a decade and the climate anomalies caused by atmospheric soot decreased proportionally, with respect to severity of the scenario. Across all scenarios, surface shortwave radiation reached its all-time low two years post-war, corresponding to the point at which *Cycles* modeled crops were simulated with the lowest average solar radiation (Fig. 2). Consequently, global surface temperature immediately and rapidly declined after the catastrophe and on average reached its lowest point in the third year post-war, with more extreme cooling in the Northern Hemisphere^6,34^. Our crop models summarized this cooling trend. Daily average temperature for landraces modeled by *Cycles* reached its lowest point two to three years post catastrophe. Barley, our crop with a primary distribution in the Northern Hemisphere, experienced the coolest post-war temperatures (Fig. 1a; Fig. 2). In the coolest year of the 150 Tg Russia-US scenario, daily average temperature of the growing season across all simulated accessions decreased by 11.3 °C for maize and sorghum, 13.1 °C for rice subsp. *japonica*, 14.3 °C for rice subsp. *indica*, and 12.1 °C for barley as compared to the averaged control daily temperature across years, indicating the severity of this scenario.

For maize, rice, and sorghum, whose landraces modeled in this study were mostly tropical, declines in temperature across the simulated growing season led to an increase in the number of days required for a plant to reach maturity. The strength of this relationship increased with the more severe soot scenarios (Fig. S1). Failure to accumulate enough thermal time during the growing season was recorded as the crop not reaching maturity. As the *Cycles* set up did not account for genetic and adaptive variation among landraces, an individual not projected to reach maturity can be interpreted as environmental conditions that are relatively inhibitory for growth. The simulated number of days to maturity generally corresponded to the severity of the climate anomaly of the post-war soot scenario. The most extreme environmental effects of the 150 Tg scenarios at least doubled the number of days to reach maturity for all tropical crops (Fig. 2). In the second post-war year of this scenario, 90% of barley, 62% of rice subsp. *indica*, 51% of rice subsp. *japonica*, 54% of maize, and 33% of sorghum accessions were projected to not reach maturity.

### Identification of environmentally adaptive genetic loci

For each crop species, we acquired published genotype data of landrace accessions used in *Cycles* simulations above for use in modeling and predicting disruptions to current genotype-environment relationships. The final set included 6,384 accessions with genotype data represented by various sequencing and genotyping methods: 215 barley accessions with exome sequencing (1,688,807 single nucleotide polymorphisms, (SNPs))^35^, 3,404 maize accessions with genotyping-by-sequencing (GBS) (946,072 SNPs)^36^, 986 rice accessions with whole genome resequencing (WGS) (677 subsp. *indica*, 309 subsp. *japonica*; 9.78 million SNPs)^22^, and 1,779 sorghum accessions with GBS (459,304 SNPs)^37^. Though differences in genotyping methods and the distribution of genotyped accessions may influence our ability to model adaptation, we sought to identify datasets that most represented the diversity of genotypes and environments that landraces of our focal species originate from and are likely adapted to.

We built gradient forest (GF) models that were used to represent current genotype-environment relationships and for predicting maladaptation in crop landraces following post-war soot induced change in climate. For use in GF models, we first identified a subset of genomic loci that we hypothesized were more likely to underlie local adaptation. Specifically, we identified genetic loci that were associated with landrace climate of origin and flowering time quantitative trait loci (QTL) for use in GF models. Following methods described in ^38^ and for each crop species, we used partial redundancy analysis (pRDA) to identify the top 1,000 genetic loci associated with variation in 13 *Cycles-*derived environmental variables under the control scenario while also accounting for population structure (methods; Fig. S2). To ensure potentially critical phenology QTL were accounted for in our models, we further identified single nucleotide polymorphisms (SNPs) of loci found within and in cis-regulatory regions (+/-5 kilobase (kb) pairs) of known flowering time network genes (Table S2). We identified loci known to be involved in flowering time for each crop species by literature review, obtained gene coordinates for each flowering time gene, extracted all SNPs that overlapped within and in cis-regulatory regions of the genomic region, and filtered each species’ set of flowering time loci to account for patterns of linkage disequilibrium. The number of flowering time SNPs included in our GF models for each of our focal species included 636 for barley, 608 for maize, 314 for rice subsp. *indica*, 323 for rice subsp. *japonica*, and 116 for sorghum (differences in number are a product of marker density). In total, the final genetic dataset used to build each species’ GF model included the top 1,000 SNPs associated with variation in the control *Cycles*-derived environmental variables identified by pRDA and the SNPs found in and near flowering time network genes.

### Control scenario GF models describe existing genome-environment associations

We built GF models representative of current genotype-environment associations using the loci described above and *Cycles*-derived environmental variables of the control simulation, averaged across all years of *Cycles*-simulated growth, for each crop species. GF is a nonparametric multivariate approach that fits an ensemble of regression trees using Random Forest^39^ and models changes in allele frequency along environmental gradients^40^. GF’s functions provide a means to rescale environmental predictors from their normal units (e.g., °C, mm) into a unit of cumulative importance for describing variation in a genetic dataset. For all GF models, the emergent environmental parameter of simulated days to maturity was in the top five most important predictors for describing variation in the genetic dataset of loci we hypothesized to contribute to environmental adaptation (Fig. S3). Across all crop species, no single environmental variable was substantially more related to turnover in allele frequencies of tested loci, indicating that GF models captured genome-wide relationships to multiple environmental gradient signals rather than a high impact at a single locus^30^. The differing importance of environmental variables specific to a growth stage of plants (variable constrained to the vegetative or reproductive stage of growth) indicated that stress experienced by plants changes across the different phenological stages of growth and genetic variation can be associated with life-stage specific stress.

### GF models capture adaptation in landraces

To test if GF models (constructed using *Cycles*-derived environmental outputs from the control scenario and the set of loci we hypothesized to be important in local adaptation) captured current environmental adaptation in landraces, we compared published performance data of 11,762 maize landraces grown across 13 common gardens in Mexico^27,36^ to predicted GF offset of genotype-environment relationships of landraces grown in common gardens. The common gardens maize landraces were grown and phenotyped in spanned geographic and environmental range of maize cultivation (Fig. S4A). We predicted how ‘adapted’ maize landraces accessions were to the common gardens they were grown in through calculation of GF offset. We calculated GF offset for each maize landrace accession grown in each common garden as the Euclidean distance between the accession’s control GF modeled genotype-environment association (representing current genotype-environment relationships) and the expected genomic composition at the common garden (representing the optimal genotype-environment relationship for a common garden). As offsets are calculated from current genotype-environment relationships in the GF model, they are weighted by the contribution of different loci that are involved in current landrace adaptation and indicate what amount of genetic change would be required for adaptation to a common garden. Accessions with a low GF offset are expected to be better adapted to the conditions at the common garden, as they require less genetic change to be adapted to the environment of a common garden. We found that, indeed, accessions performed best (height and yield measures) when grown in sites where they had low GF offset (Fig. 3A; Fig. S4B). Furthermore, anthesis silking interval (ASI, synchronicity of male and female flower maturity) was reduced when accessions were grown at sites in which they had lower GF offset. ASI is a reliable predictor of stress in maize^41^, indicating maize landraces were less stressed when grown in common gardens to which they were predicted to be adapted (lower GF offset).

**Figure 3.**
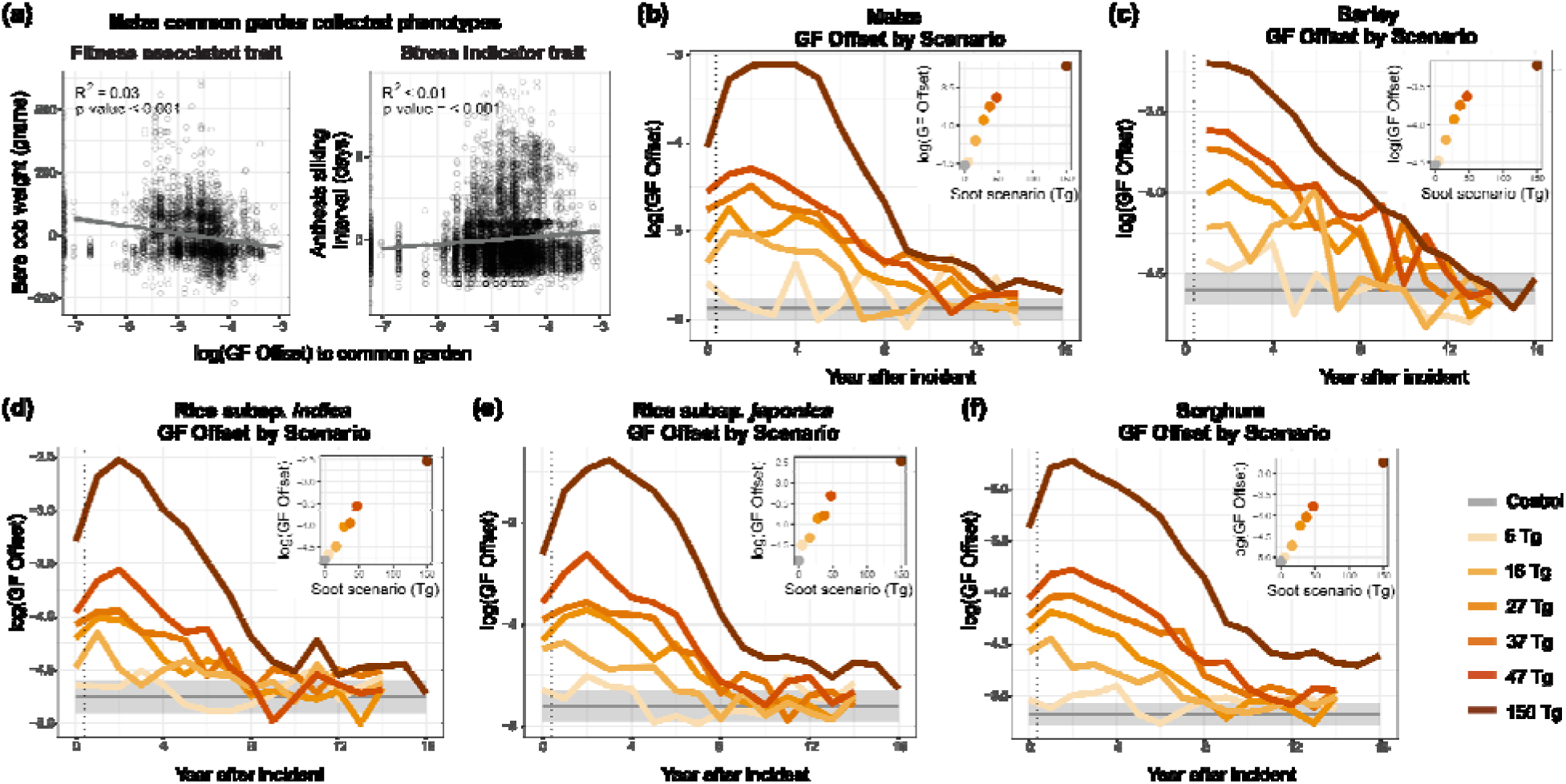
GF models capture current genotype-environment associations in maize landraces and were used to predict maladaptation (GF offset) under post-catastrophic climate scenarios. A) Phenotypic residuals (remaining variation after accounting for experimental design) plotted against the logged GF offset of maize landrace accessions grown in common gardens. GF offset is calculated for each phenotyped accession as the Euclidean distance of the expected genotype-environment relationship at a common garden common vs the genotype-environment relationship from the accessions’ point of origin. Points with a more negative logged GF offset indicate maize landrace accessions that are expected to be adapted to the common garden they were grown in. Yearly logged GF offset for B) maize C) barley D) rice subsp. *indica* E) rice subsp*. japonica* and F) sorghum. The control line shows the mean logged GF offset across all years, with the shaded region representing the standard deviation of yearly means to indicate fluctuations in maladaptation (GF offset) due to normal variability in climate. Soot scenario lines are averaged logged GF offset across all accessions of a species and colored by soot scenario. The vertical dotted line indicates the time of soot injection into the climate models. Inlaid scatter plots are the averaged logged GF offset across all accessions of a species two years after the incident.

### The degree of GF offset post-catastrophe follows the magnitude of climate disruption

After confirming the fitted control GF models captured adaptation in landraces, we then used GF models to predict the expected locally-adapted genomic composition for landraces across space and time under the six post-war scenarios. To predict the magnitude of maladaptation, we calculated GF offset as the Euclidean distance between a given landrace source location’s expected genomic composition between control (representing current genotype-environment relationships) and the six soot scenarios (representing the optimal genotype-environment relationship for a soot scenario) separately. High GF offset values corresponded to a greater degree of maladaptation and represented a greater shift in allelic composition required for adaptation to persist in the climate produced by the soot scenario. For all crops and scenarios, GF offset values followed the trend in post-war climate disruptions, with a sharp increase and gradual recovery after 10 to 15 years (Fig. 3B-F; Fig. S5). GF offset for all crops reached its highest point two to three years post-catastrophe, indicating that crops were expected to have the highest degree of maladaptation when global solar radiation and temperatures reached their all-time low. Maximum GF offset of each target scenario linearly corresponded to the amount of soot simulated for the 5 Tg to 47 Tg soot scenarios. In the most extreme 150 Tg scenario, the trend was more pronounced and deviated from the linear pattern (Fig. 3 B-F). Across all crop species, we detected a strong latitudinal pattern associated with GF offset values, equatorial regions which experienced less adverse climate impacts were predicted to be less maladapted to post-war conditions (Fig. 4).

**Figure 4.**
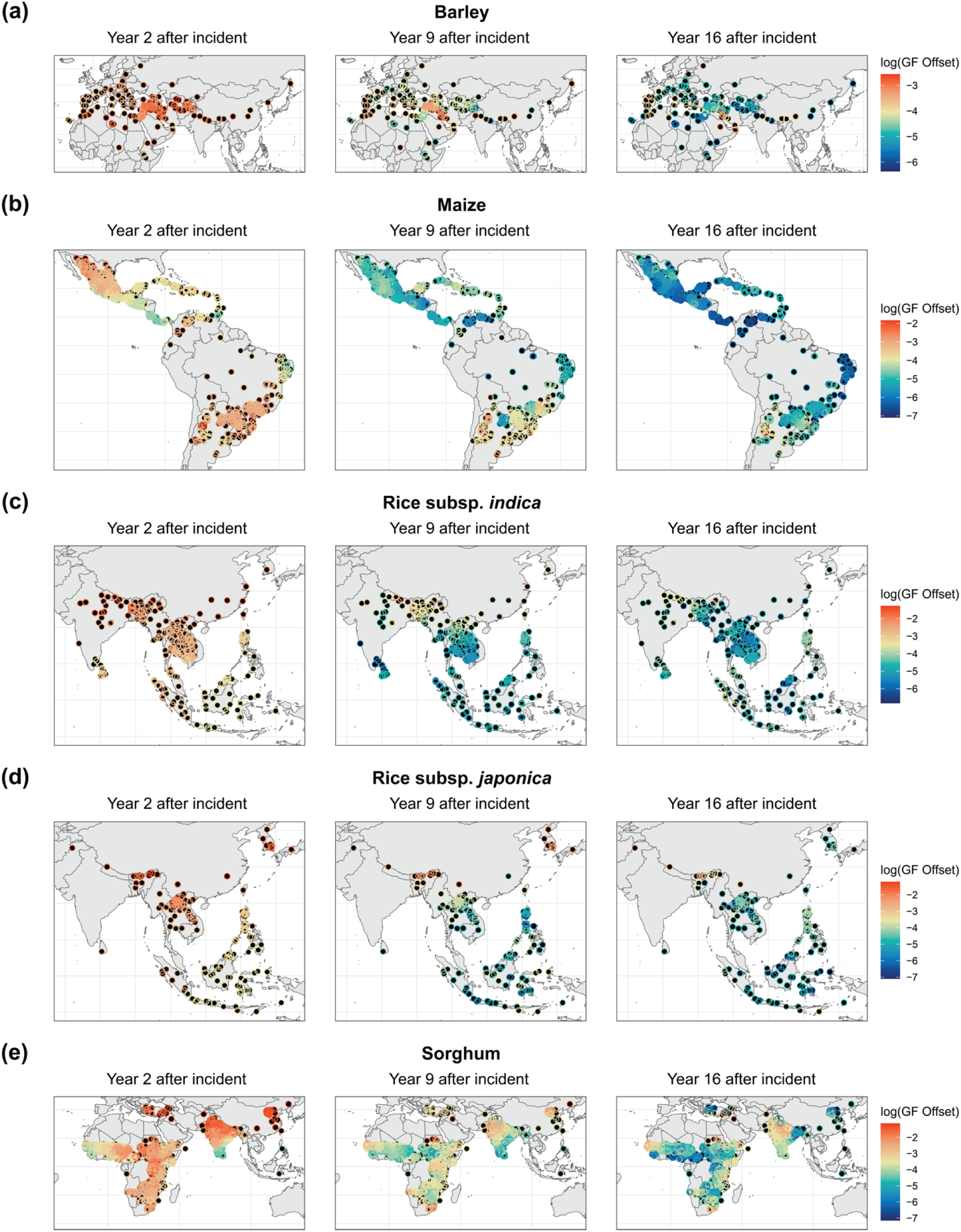
Global distribution of logged GF offset (predicted maladaptation) under the 150 Tg scenario 2, 9, and 16 years after the incident for landraces (filled black circles, with overlaid open colored circles indicating offset). Higher GF offset values correspond to a larger degree of predicted maladaptation under the post-war scenario.

### Identification of landrace substitutions for post-catastrophe adaptation

We next leveraged our GF models to identify landrace genotypes best matched to specific post-catastrophic conditions, indicating potential varietal substitutions for locations with landraces that were the most maladapted to post-catastrophic climates. Under post-catastrophic conditions, many locations will not have climate suitable for the cultivation of crops and we constrained our analyses to only look for substitutions for locations that were projected to have a crop reach maturity in the worst year (year 2 post-strike) of the 150 Tg scenario. After filtering for locations that were not expected to be suitable for agriculture, 10% of barley, 38% of rice subsp. *indica*, 49% of rice subsp. *japonica*, 46% of maize, and 67% of sorghum landraces source locations were retained to search for a suitable substitution. For the remaining locations, we identified the most vulnerable locations as those with the highest GF offset (predicted maladaptation). We then searched for the most optimal substitution globally as well as the best within country substitution, identifying the landrace accession with the lowest GF offset to the post-catastrophic climate in the vulnerable location (Fig. 5). Though the identification of landraces with lower levels of maladaptation to post-catastrophic conditions may be valuable for finding the genotypes most resilient to post-catastrophic climates, it is important to note that our calculation of maladaptation is a relative metric and to approach these findings with caution. The post-catastrophic climate of the 150 Tg scenario may be sufficiently extreme to dramatically reduce the absolute production of accessions that is identified as a suitable substitution and predicted to have a low GF offset to the novel climate conditions.

**Figure 5.**
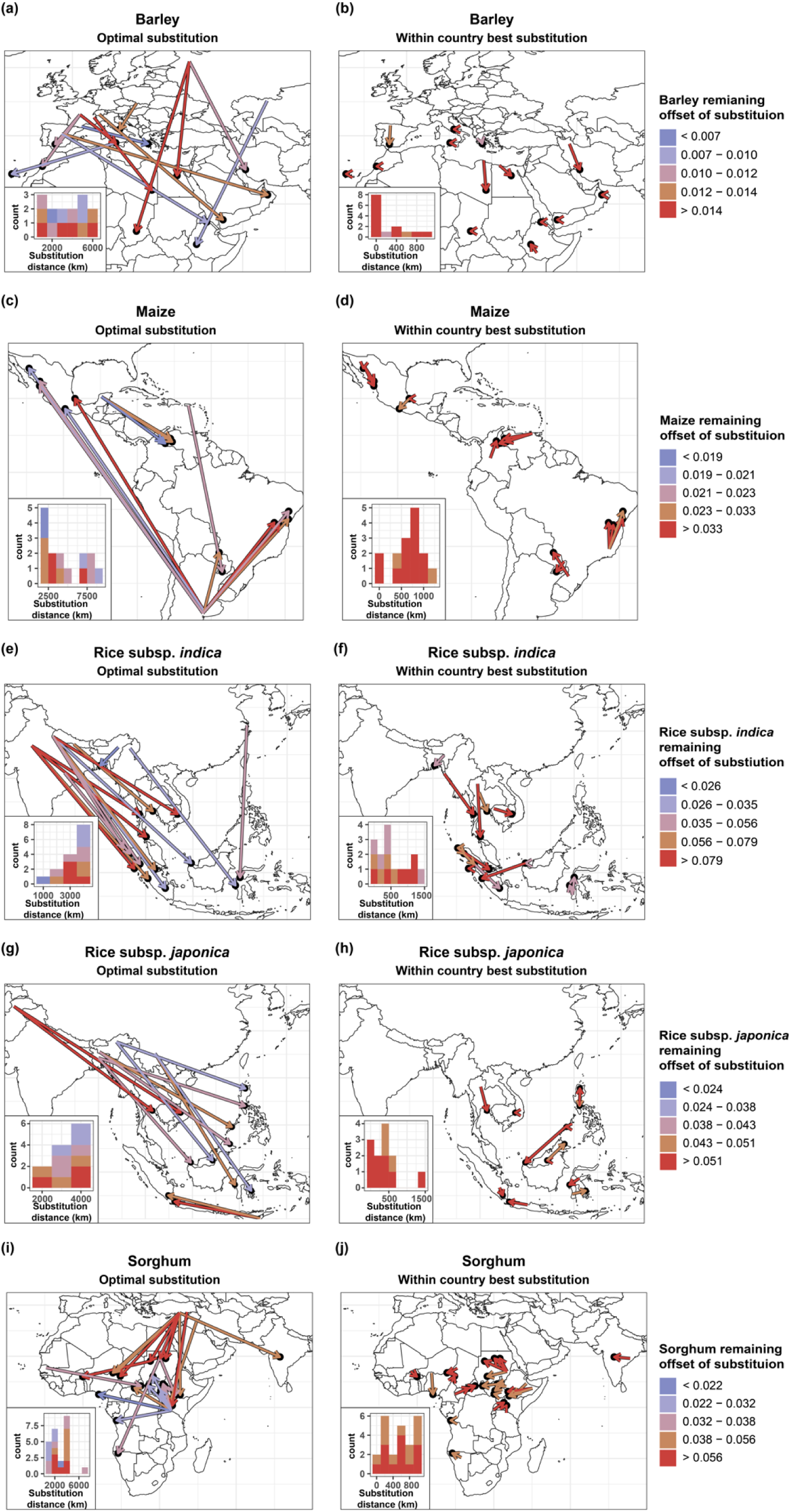
Substitution trajectories for the most vulnerable landrace populations in year 2 of the 150 Tg scenario. For each crop, arrows connect the source location of a landrace accession that is the most optimal to the vulnerable location (arrowhead) and are colored by the remaining GF offset (maladaptation) of the substitution. Substitutions are colored by how well-matched the moved landrace is to the vulnerable location, where colors corresponding to lower GF offset of substitution indicate a substitution that has a low degree of maladaptation to the novel environment. For each crop, substitution trajectories are provided for the most optimal substitution across all available germplasm and the best within-country substitution. Inlaid histograms represent the frequency of substitutions of different distances and are colored by the remaining GF offset (maladaptation) of the substitution.

Across all crops, the most optimal substitution was often far away (∼1000 to ∼10,000 km) and across country borders. For many locations, the best substitution still had a high degree of GF offset, indicating that there was not a genotype that was expected to be adapted to the post-catastrophic climate at the vulnerable location included in our dataset (Fig. 5A, C, E, G, I). This could be due to the severity of the novel climate at the vulnerable location, the absence of a landrace accession that was expected to be adapted to the novel environment, or some combination of both. For all crops, the best substitution trajectories typically moved landrace accessions from poles and high elevations towards the equator and low elevations, indicating that landrace germplasm adapted to currently cooler climates may be sources of resilience for locations that may be more likely to support agriculture post-catastrophe. For all crop species, there were instances where one genotype was the most optimal substitution for multiple vulnerable locations, suggesting genotypes that may be particularly valuable for post-catastrophic agriculture.

In the case of a catastrophe, substitutions across long distances may not be possible due to socioeconomic disruptions, e.g. in transport and trade. We further searched for the optimal within-country substitution. For all crops, within country substitutions with a low GF offset were rare; within country substitutions always had a higher GF offset, corresponding to higher expected maladaptation, than the optimal global substitution (Fig. 5B, D, F, H, J). Though maintaining a high degree of maladaptation (GF offset), most within-country substitutions included trajectories moving individuals towards the equator and lower elevations.

The within-country current diversity of environments to which landraces are adapted may be important for finding a suitable substitution. To test this hypothesis, we compared the GF offset for the 25% most maladapted locations within each country after using global substitutions versus within-country substitutions. We focused on sorghum because it was the crop with the most countries having viable cultivation in year 2 of the 150 Tg scenario, giving power to compare countries. As expected, all 31 countries with at least 5 sorghum accessions had greater GF offset for the most maladapted locations when only using within-country substitutions, compared to the global substitutions. The proportional inferiority of within-country compared to global substitutions was only weakly related to the number of landraces from each country (r = 0.27, p = 0.14). We next tested if the control climate mean and variance influenced the inferiority of within-country substitutions in a multiple regression, while accounting for the number of landraces in each country. We found that the countries with less variance among landraces in cold stress and greater mean cold stress had significantly worse within-country substitutions compared to global (linear model, mean cold t *=* 3.99 p = 0.0005, variance in cold t = 2.54 p = 0.0173, number of landraces t = 0.37 p = 0.7173, R^2^ = 0.45). This highlights the potential future value of diversity for regions and nations housing landraces adapted to diverse climates.

## Discussion

The resilience of agricultural systems to changing climate determines global food security. In this study we used information on landrace genetic variation and environment of origin for agronomically important cereal crops to predict disruptions to their adaptation/cultivation and to explore if the diversity of landraces may be beneficial sources of resilience in the case of a soot-producing climate catastrophe. Consistent with other groups who have investigated the consequences of a soot-producing catastrophe on global agriculture^7,9^ and fisheries^34^, we find the climate impacts would be devastating to global subsistence agriculture, many locations would become unsuitable for agriculture, and for the most extreme soot scenario, the locations that remain suitable may not have sufficient local landrace diversity within a species to enable a successful substitution of a resilient variety.

Our crop model results correspond to previous estimates of the climate impacts of soot-producing catastrophes^6,7,9,32,34,42^ while also providing an assessment of the diversity of environments to which crop landraces of globally important cereal crops are adapted. Increases in the number of days simulated to reach maturity corresponded to the climate anomalies of reduced daily temperature and solar radiation. In the years and locations with the greatest climate impacts, landraces in higher latitudes rarely achieved full maturity. Colder temperatures slow down phenological development, and can diminish photosynthetic activity and damage tissue.

We built GF models to summarize current landrace genotype-environment relationships and validated that GF models captured real adaptive differences through use of phenotypic data collected for a broad diversity panel of maize landraces grown in common gardens across Mexico. We show that predicted maladaptation, in the form of GF offset, is associated with height, yield, and stress-related traits, demonstrating a new test of these tools^23^. Landrace accessions had classic phenotypic patterns of local adaptation when grown in common gardens they had low GF offset (maladaptation) to, suggesting GF models captured broad adaptation of landraces’ local environments^43^. However, landrace performance was not perfectly predicted by our genotype-environment model. This inability to completely predict adaptation may be attributable to limitations of genotype-environment association approaches or to the maintenance of diversity within environments. In general, reciprocal transplants and common gardens often find mixed evidence for local adaptation^44^. Genotypes from the same environment may differ in performance in a given common garden environment if processes like migration or environmental fluctuations maintain diversity within populations or if important selective forces are not present in experimental conditions. Our validation methodology confirms that GF offset can be a powerful tool to capture current genotype-environment relationships though our inability to perfectly predict adaptation likely highlights a potential importance of maintaining genetic diversity within a site, which may complicate our ability to model these relationships.

When environments change and populations are not able to track the environmental change through plasticity or rapid shifts in genetic composition, populations may become maladapted and have reduced fitness in a novel climate^45^. In our case, landraces were predicted to be the most maladapted, or have the highest GF offset, in the locations where climate was the most disrupted from long-term averages, corresponding to the most extreme soot scenarios and the years post-war where atmospheric soot was the most abundant. The strong relationship we observed between GF offset and soot-induced change in climate is perhaps unsurprising. GF models are trained using current genotype-environment associations and any shift in the environment will likely require a change in genomic composition to track adaptation to a novel climate. The ability to interpret the magnitude of offsets derived from GF-derived functions in an ecologically meaningful way has recently become a point of discussion. Genetic-based quantifications of adaptation^38^ and offset^46^ can be biased for unsampled areas or if the projected environment exceeds what is used to train the model. Though we have a broad sampling of landrace accessions for each focal crop species that are adapted to a diversity of environments and used in GF genotype-environment models, the extremeness and novelty of post-war climates used in this study likely make predicting maladaptation difficult^47^. At the same time, though the true magnitude of maladaptation may be difficult to quantify, our GF models allow us to incorporate measures of climate-associated genomic variation for the identification of the most vulnerable locations that will likely require a varietal substitution. Additionally, our GF models provide insights to the aspects of the environment that may be most related to a crop’s current adaptation, which is likely related to the evolutionary history and cultivation practices of the crop. For example, GF identified average temperature and solar radiation experienced in the reproductive growth phase as most related to rice subsp. *indica* genome-wide allelic turnover, suggesting these variables may be important in driving local adaptation within this species. Rice landraces of the *indica* variety are traditionally cultivated in warm, tropical to subtropical locations and may have limited cold tolerance^48^. While cold and solar radiation are the variables most altered by nuclear winter, perhaps suggesting vulnerability of this species, the GF model also suggests that indica genotypes vary in their adaptation to temperature and light, suggesting there is some mitigation possible with genotype substitutions.

Crop diversity has been suggested as a potential solution to mitigate climate impacts on agriculture^24,49^. For all crops included in this study, we found that landraces accessions with a distribution farther from the equator were most maladapted to post-catastrophic climates and were most often selected as the best varieties for substitutions. Most substitutions that were well matched to vulnerable locations required long migration distances and for many locations, a landrace adapted to the novel environment at the vulnerable location does not exist within our dataset. Substitutions that maintained a high level of GF offset indicated landrace varieties that may remain maladapted to the novel climate, and no other varieties were better adapted to the vulnerable, tested location. At the same time, for locations where the cultivation of crops remains possible, the identification of multiple suitable genotypes may be important for the maintenance of crop diversity within a site. For smallholders, the development of elite farmer-preferred varieties and the introgression of alleles adapted to novel climates is a priority^50^, and genotype substitutions identified here could be potential donors of such alleles. For vulnerable locations that were not predicted to have a well-adapted substitution, switching cultivation to faster-maturing crop varieties, or other non-cereal crop species that tolerate lower temperatures (e.g. potato)^51^, may be a strategy for increased resilience. However, the adoption of a new crop species requires a significant investment by farmers and substantial modifications of farmer and consumer behavior^52^. It is worth noting that there may be some diversity in response to post-catastrophic conditions in modern elite crop varieties cultivated in wealthier nations, which are not accounted for in this study. Other studies have considered changes to global crop productivity under nuclear conflict, including Jagermeyer et al. (2020) who showed that even a relatively small nuclear strike (e.g., 5 Tg of soot) would drastically impact crop production.

Though our study highlights maladaptation in cereal crop landraces following a soot-producing catastrophe, methodology used in this study can also be leveraged to understand disruptions to adaptation and possible genotype substitutions (also known as assisted gene flow) given any change in climate, including greenhouse gas induced climate change^53^. Our results indicate that for the landrace populations most vulnerable to a climate catastrophe, the within-species genetic diversity in a country may not be sufficient for resilience and substitutions across country borders of further distances may be required.

## Methods

We used landraces to characterize global disruptions to adaptation and identify resilient accessions in the case of a climate catastrophe that produces soot. Selected landrace crop species fulfilled two criteria - 1. Landrace relatives of the species account for a large portion of accessions currently grown and 2. High quality sequencing data of geographically diverse accessions were publicly available. From these criteria, we selected four cereal crop species-*Hordeum vulgare* L. (barley), *Oryza sativa* L. (rice) subsp. *indica* and *japonica*, *Zea mays* L. (maize), and *Sorghum bicolor* (L.) Moench (sorghum). For all analyses, the rice subsp. *indica* and *japonica* were run separately. Altogether, the species cover most of the agricultural areas of the globe and are cultivated in and adapted to diverse climate regimes.

### Weather data

Previously published weather data described in Toon et al. (2019) and Coupe et al (2019) simulate the climate impacts of India-Pakistan and US-Russia wars using the Community Earth System Model (CESM, version 1.3) with the Whole Atmosphere Community Climate Model Version 4 (WACCM4, version 4) as the atmospheric component, or CESM-WACCM4^54^. To more accurately represent the evolution of smoke injection, the Community Aerosol and Radiation Model for Atmospheres (CARMA;^55,56^) is coupled with WACCM to simulate the injection, lofting, advection, and removal of soot aerosols^42,57^.

The climate impacts of nuclear war were simulated by injecting varying quantities of black carbon aerosol (soot) into the stratosphere in a layer between 100 and 300 hPa over a 1-week period starting on 15 May above the U.S. and Russia, or the South Asian subcontinent^6,32,42^. In total, six nuclear war scenarios were simulated, and we refer to the year soot was injected as year “0”. For the five India-Pakistan nuclear war scenarios (soot injections of 5 Tg, 16 Tg, 27.3 Tg, 37 Tg, and 46.8 Tg, representing a range of arsenal sizes) simulations were each run for 19 years. One United States-Russia scenario with a 150 Tg soot injection was also considered, and the simulation was run for 21 years. This scenario assumes both countries use most of their nuclear arsenals^58^ and is still possible given modern nuclear arsenals. Additionally, a single control run that repeats the climate forcing of 2000 was simulated for 20 years to represent normal atmospheric circulations^6,32^.

### Cycles

The *Cycles* agroecosystem model was used to infer growth and stress variables of landrace accessions’ point of origin using conditions accessions are expected to be adapted to (control scenario) and post-catastrophe (six post-nuclear war soot scenarios). *Cycles* is a process-based multi-year and multi-species agroecosystem model^2,33^ that requires a number of input files to simulate crop growth. All simulations were carried out using *Cycles* v0.13.0 (https://github.com/PSUmodeling/Cycles). The crop description file defines the physiological and management parameters that control the growth and harvest of crops used in the simulation. For each of our crop species, we used *Cycles* default crop parameters from the default crop description file. The management (operation) file defines the daily management operations to be used in a simulated crop rotation. We activated conditional planting where *Cycles* “plants” a simulated crop once certain soil moisture and temperature levels are satisfied within a window of planting dates. For many of the scenarios where planting conditions are not met (i.e. daily temperature remains too low) *Cycles* forced planting on the last day of the planting window. We turned on the automatic nitrogen fertilization option and set planting density to 67% for all crops in the simulation to be grown without nitrogen limitations so that stress observed in model outputs was due entirely to climatic factors. Weather files were built using the CESM-WACCM4 outputs as described in ^32^ and ^6^ for one control and six post-nuclear soot scenarios, formatted for use in *Cycles*. The weather files were generated by aggregating the three-hourly CESM output to daily time steps at all CESM grids, which have a 1.9° latitude × 2.5° longitude resolution. Weather files were matched to landrace point of origin for each simulated accession, where the climatic parameters used to simulate growth match the location accessions were sourced from. Weather files included variables describing variation in daily precipitation, temperature, solar radiation, humidity, and wind. Soil physical parameters were obtained from the ISRIC SoilGrids global database^59^ via the HydroTerre data system^60–62^ for all simulation locations. Soil files were also matched to landrace point of origin for each simulated accession and describe the average soil characteristics and land use for crop cultivation types. For accessions designated as paddy rice by ^22^ we used the irrigated or post-flooding land use type. Rainfed land use type was used for all other simulated crop accessions.

For all simulated accessions of each crop species, seven *Cycles* simulations, including the control scenario and six soot scenarios were implemented separately. *Cycles* models simulated 20 years of crop growth for the control scenario, 15 years of crop growth after impact for the India-Pakistan scenarios (5 Tg, 16 Tg, 27 Tg, 37 Tg, 47 Tg, and 150 Tg), and 17 years of crop growth for the US-Russia scenario (150 Tg). From the outputs of each *Cycles* simulation and for each year growth was simulated, we extracted variables summarizing the environmental stress and simulated growth plants experienced for each of our focal crop species (*Cycles*-derived environmental variables). Variables included information on the number of days to reach maturity, water stress, cold stress, and light stress experienced across simulated plant growth and when in the vegetative and reproductive phase (Table S1). For accessions not projected to reach maturity, certain environmental summary variables were not extractable, and we imputed the 95% stress of the variable for each accession with missing environmental values, specific to crop species and the year growth was simulated for.

### Genotyped datasets

As differences in genotyping resolution across species might influence the detection of genomic signals of adaptation, we selected datasets with high density genomic markers and a distribution of sequenced landraces accessions that most represented the environments that landraces of our focal species originate from and are likely adapted to. Advances in technology have made low-coverage whole-genome sequencing (WGS) relatively inexpensive, providing datasets that are particularly well-suited for research exploring polygenic signals. All genotype files were processed in PLINK, an established software for analyzing and filtering genotypic data^37^. For each landrace species, raw genotype files were filtered for minor allele frequency (MAF) removing all SNPs with lower than 5% MAF and for linkage disequilibrium (LD) to reduce the number of SNP candidates we tested for environmental association. As the initial genotype files differed in size, the LD filter step included different conditions to thin files. We used --indep-pairwise 30 10 .1 for both rice subsp. (*indica* and *japonica*) and sorghum, and --indep-pairwise 100 10 .05 for the maize and barley data files. This filtering step resulted in 74,430 SNPs for barley, 43,818 SNPs for rice subsp. *japonica*, 61,430 SNPs for rice subsp. *indica*, 67,522 SNPs for maize, and 20,387 SNPs for sorghum to test for association to the species-specific *Cycles*-derived environmental variables.

### Genome scan for environmentally related SNPs

Genotype-environment associations test for genetic variation that is statistically correlated to environmental predictors. We followed partial redundancy analysis (pRDA) methods developed by ^38^ to identify loci putatively involved in environmental selection for our focal crop species. For each crop species, pRDA models were built using population allele frequencies (population defined as accessions from the same geocoordinates) from the filtered genetic dataset as response variables and the 13 *Cycles*-derived environmental variables from the control simulation, averaged across the 20 years of modeled growth as explanatory variables. Neutral genetic structure was accounted for by including the first three axes of a population PCA as conditional covariables. Using the *rdapat* function described in ^63^, we identified the top environmentally related (outlier) loci based on the extremeness of their loading along a Mahalanobis distance distribution calculated between each marker and the center of the first two pRDA axes. P-values for each marker were derived as this distance, corrected for the inflation factor using a chi-squared distribution with two degrees of freedom. We then selected the top 1,000 markers with the lowest P-values as candidate outliers to represent loci that may be important for environmental adaptation. The analysis was carried out using R/vegan^64^.

To assess whether the top loci selected by pRDA are unique to the method, we further implemented Bayesian-information and Linkage-disequilibrium Iteratively Nested Keyway (BLINK) and compared the significant loci as identified by BLINK and pRDA for sorghum. BLINK is a package commonly used for genome-wide association studies (GWAS) and improves upon traditional GWAS methods by addressing limitations such as computational inefficiency and reduced statistical power^65^. We ran BLINK separately for the same 13 *Cycles*-derived environmental variables used in the sorghum pRDA model and extracted the set of significant loci (p-value < 0.05) for each BLINK model that were built separately for each climate variable. For all BLINK models, the first three axes of a population PCA were used as covariates to account for population structure. We then compared the set of BLINK-identified significant loci across all 13 models (8,728 unique SNPs) to the 1,000 most significant loci as identified by pRDA and found that 556 SNPs were present in both datasets. Thus, the overlap between genotype-environment association methods for identifying loci that are related to variation in environmental gradients confirm that the results are not highly sensitive to the approach.

### Identification of flowering time SNPs

We further accounted for genetic variation that may capture important plant phenological processes by including SNPs of known flowering time network loci for each focal crop species. We conducted a literature search to identify genes known to be involved in the flowering time network for each crop (Table S2). Gene coordinates of each flowering time gene were gathered from the gff3 files that corresponded to each reference genome used to call SNPs (maize (reference B73v2, https://figshare.com/articles/dataset/GTF_and_GFF_for_maize/895628); rice (reference R498 IGDBv3, http://mbkbase.org/R498/); sorghum (reference S. bicolorv3.1, https://phytozome-next.jgi.doe.gov/info/Sbicolor_v3_1_1)^66–68^. For maize, rice subsp. *indica* and *japonica*, and sorghum we also included SNPs found +/-5 kilobase (kb) of each flowering time gene to account for variation in cis-regulatory elements. Barley sequence information was reported as contigs and we extracted SNPs located in contigs previously identified to overlap with homologs of well-characterized genes in *Arabidopsis thaliana*^35^. Gene coordinates for the location of each flowering time gene region or flowering time related contig extracted using --extract in PLINK^69^. To account for patterns of linkage disequilibrium, we further filtered each species’ set of flowering time loci (gene and sites up and downstream of the gene) and only retained SNPs with an r² value less than 0.2 within the flowering-time genic window and flanking region.

### Gradient forest models and calculation of offset

Gradient forest (GF) is a machine learning algorithm extended from random forest which searches for genotypic patterns as associated with environmental descriptors. Using R/gradientForest::gradientForest^40^, we built GF models to associate current adaptive allelic diversity (the combined set of pRDA-identified environmentally related loci and flowering time network loci) with *Cycles*-derived environmental variables from the control simulation, averaged across the 20 years of modeled growth (hereafter, control GF model). Models were built separately for each of our focal crop species to describe control species-specific genotype-environment relationships. The control GF model parameters were tuned to increase the number of trees built to *ntree* = 500.

Genomic offset (also known as genomic vulnerability) is one metric used to characterize maladaptation with a genomic context (reviewed in ^28^). The distance between current and expected genotype-environment associations under some change in environment is representative of the genomic offset, or the genetic shift required in a population to adapt to the future climate. Comparing the control genotype-environment association captured by GF models (control GF model), and the projected genotype-environment association for different scenarios (common garden, target scenarios, vulnerable locations) we made several measurements of GF offset to summarize predicted maladaptation. For all GF offset calculations, we followed methods described in ^24^.

To validate that our control GF models captured current genotype-environment associations, we first used the maize control GF model to predict the genomic composition expected at common gardens maize landraces had been phenotyped in for a previous study^36^. Here, the GF offset was defined as the Euclidean distance of current genotype-environment relationships at the common garden site from the genotype-environment relationship of each landrace’s point of origin. This measure summarized how genetically well matched a landrace was to the common garden it was grown in (measure of predicted maladaptation to a common garden) and was compared to the phenotypic breeding values for each landrace grown in a common garden.

### Validation of gradient forest-predicted adaptation

We used phenotypic data of maize landraces grown in 23 trials across 13 common garden locations over 2 years to confirm that our control GF models captured real differences in current landrace genotype-environment relationships. We restricted our analysis to include the phenotypic data of landraces accessions that were simulated in *Cycles* models.

Briefly, phenotyped accessions are a part of the broader SeeD evaluation of the maize landrace collection^36^. Accessions were planted in multiple environments under a replicated F1 crossing. Importantly, two features of the crossing design ensure that phenotype data is not overly biased by elevational adaptation. Crossed plants were preferentially grown in locations that were of similar adaptation (highland tropical, sub-tropical or lowland tropical) to their home environment and each plant was crossed to a tester that was adapted to the environment that the F1 seeds were grown in. These design features allowed for comparison of a larger sample of accessions, but also led to an unbalanced experimental design. As a further consequence of the experimental design, apparent adaptive differences among landraces may be reduced and make phenotypic estimates of adaptation more conservative^27^. We extracted phenotypic information capturing differences in plant height (PH), the total weight of ears (kernels and cob) measured in the field (field weight; FW), bare cob weight (BCW), moisture adjusted grain weight per hectare (GWH), days to anthesis (DA), days to silking (DS), and anthesis-silking interval (ASI) for plants grown in trials (https://data.cimmyt.org/dataset.xhtml?persistentId=hdl:11529/10548233). The phenotypic datasets ranged from having n = 4,851 (BCW) to n = 11,762 (ASI) across the field sites. Following methods from Gates et al., (2019) and Romero Navarro et al., (2017), we estimated breeding values controlling for tester, checks, and field position in a complete nested model. We further accounted for the random effect of tester and year.

### Calculation of offset under post-catastrophic conditions

Once we confirmed that our maize control GF model captured phenotypic differences representative of adaptation to common gardens with conditions most like the source locations landraces are sourced from, we extended our control GF models to predict maladaptation in crop landraces under the six post-war target climate scenarios. Here, GF offset was calculated as the Euclidean distance between the current predicted genotype-environment association (control GF model) and for each soot scenation, the future projected genotype-environment association across all 13 *Cycles*-derived environmental variables.

We confirmed that GF offset values were correlated to relative changes in environment that were most related to genotype-environment associations, as summarized by the control GF model, by comparing the difference in GF offset (target subtracted from control) versus the change in *Cycles*-derived environmental variables (Fig. S6). Environmental variables were scaled and adjusted by their relative contribution to GF models.

### Identification of landrace substitutions for post-catastrophe adaptation

To understand if existing landrace diversity may be a source of resilience following a climate catastrophe, we used our GF models to identify the best suited substitutions for locations with the landraces that are the most maladapted^24,29^ under the most extreme target scenario (150 Tg) and for the year where climate is the most disrupted (year 2 after soot injection). We first excluded all locations not predicted to reach maturity, so as not to identify substitutions to locations where agriculture would not likely be possible. For the remaining locations, we defined the most vulnerable locations as those with the highest GF offset to search for both the most optimal and the best within-country substitution. Clusters of vulnerable pixels were identified using R/DBscan::dbscan^70^, which groups pixels based off proximity. Clustering was based on the geographic distance between vulnerable pixels measured with R/geosphere::distm^71^. Only clusters separated by <1000 km were retained for further analysis.

For each vulnerable cluster, a GF offset (i.e., Euclidean distance) was calculated between the projected genotype-environment association of the vulnerable location under the 150 Tg target scenario and the control GF modeled genotype-environment association (control GF model) across all landrace accessions included in the model. The lowest GF offset was defined as the minimum Euclidean distance and identified the current landrace accession predicted to be best adapted to the future climate conditions of the vulnerable area. For landrace accessions that were not perfectly adapted to the locations they were substituted to (i.e GF offset is higher than 0), this measure represented the genomic gap that still needs to be filled for the migrated varieties to be fully adapted to the conditions of their new location (assuming current genotype-environment relationships are representative of perfect adaptation). High GF offset indicated substitutions where accessions are not predicted to be well adapted to the locations they were substituted to, and no other landraces included in the model were better adapted to the projected climate of the vulnerable area.

## Data availability

The Community Earth System Model is freely available from the NCAR but requires registration at www.cesm.ucar.edu/models/cesm1.2. Additionally, atmospheric model output for the 150 Tg case (Coupe et al., 2019) is available at https://doi.org/10.6084/m9.figshare.7742735.v1. The full model outputs for all simulations are very large and stored on the PetaLibrary at the University of Colorado, which is not available to the public. However, additional data from these runs can be provided upon request.

## Code Availability

The scripts for the bioinformatics analysis are publicly available in GitHub at GitHub Repo.

## Acknowledgments and funding sources

We thank Liana Burghardt and Estelle Couradeau for comments on earlier versions of this manuscript and the whole Penn State Food Resiliency team for their input and discussion. We would like to thank Daniel Runcie and Jeffrey Ross-Ibarra for helpful feedback on analyses using the common garden data. This research was supported by the Food Resilience in the Face of Catastrophic Global Events grant funded by Open Philanthropy. This work was also supported by NIH award R35 GM138300 to J.R.L. A.K. and Y.S. were additionally supported by the USDA National Institute of Food and Agriculture and Hatch Appropriations under Project #PEN05001 (Accession 7007612) and Project #PEN05016 (Accession 7007513), respectively.

## Author contributions

C.M.M., Y.S., A.K., J.R.L designed research; C.M.M. performed research; Y.S., A.K. contributed new analytic tools; C.M.M. and J.R.L. analyzed data; C.M.M. wrote the manuscript, with input from Y.S., A.K., R.J.H.S., and J.R.L.; All authors contributed to manuscript revision.

---

**Figure S1.**
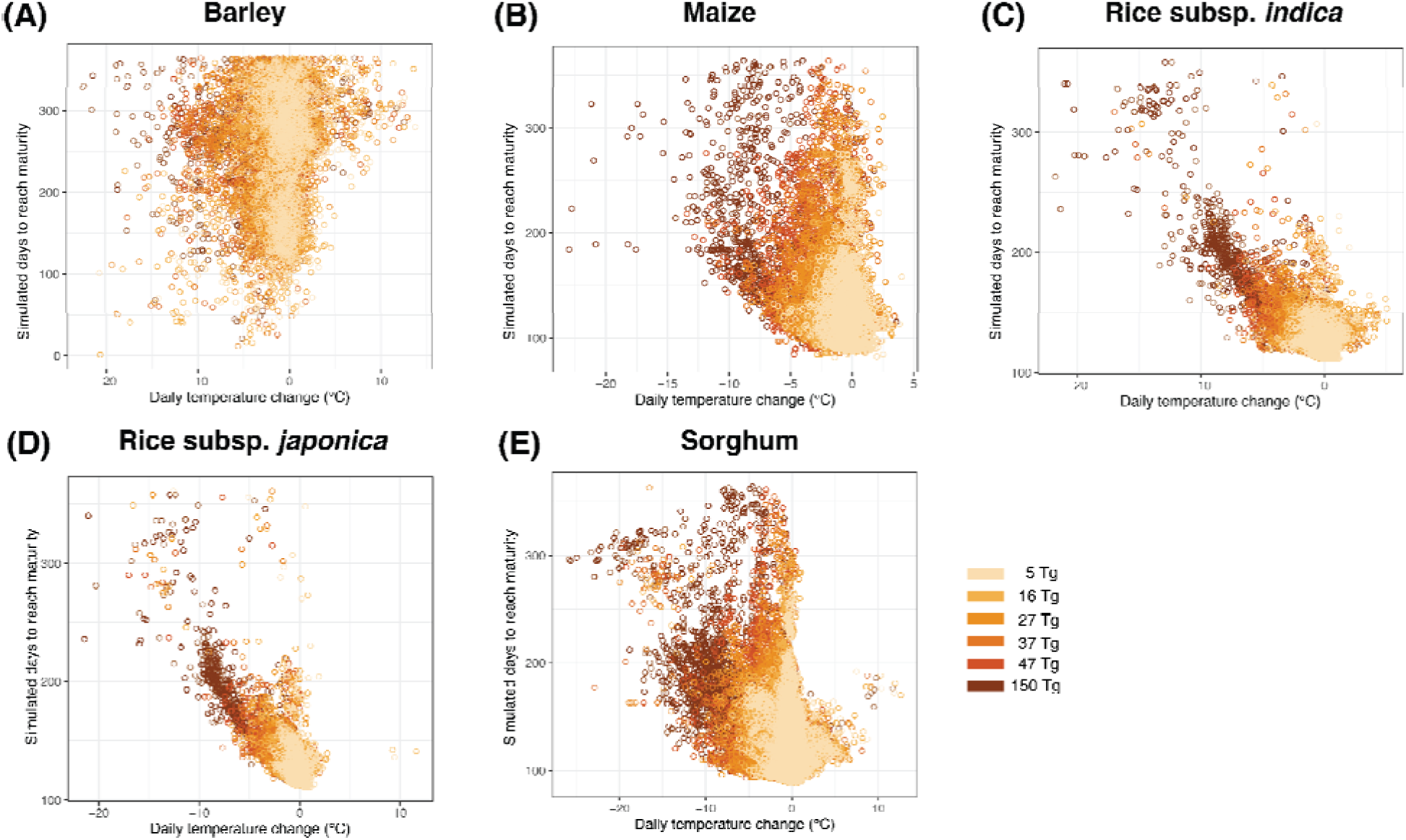
The relationship between simulated days to reach maturity and change in daily temperature for all landrace accessions across all years simulated, colored by soot scenario. Points are masked from the plots for years where landrace accessions were not projected to reach maturity. The frequency of crop failure (points that are masked from the plot) for the year with the most extreme climate impacts are given here as percentages for the 5 Tg, 6 Tg, 27 Tg, 37 Tg, 47 Tg, 150 Tg scenarios. Barley: 6%, 8%, 13%, 22%, 24%, 90%. Maize: 0%, 0%, 0%, 0.5%, 4%, 54%. Rice subsp. *indica*: 0.5%, 1%, 2%, 3%, 5%, 62%. Rice subsp. *japonica*: 3%, 5%, 12%, 19%, 20%, 51%. Sorghum: 0, 0.5%, 0.5%, 2%, 4%, 33%.

**Figure S2.**
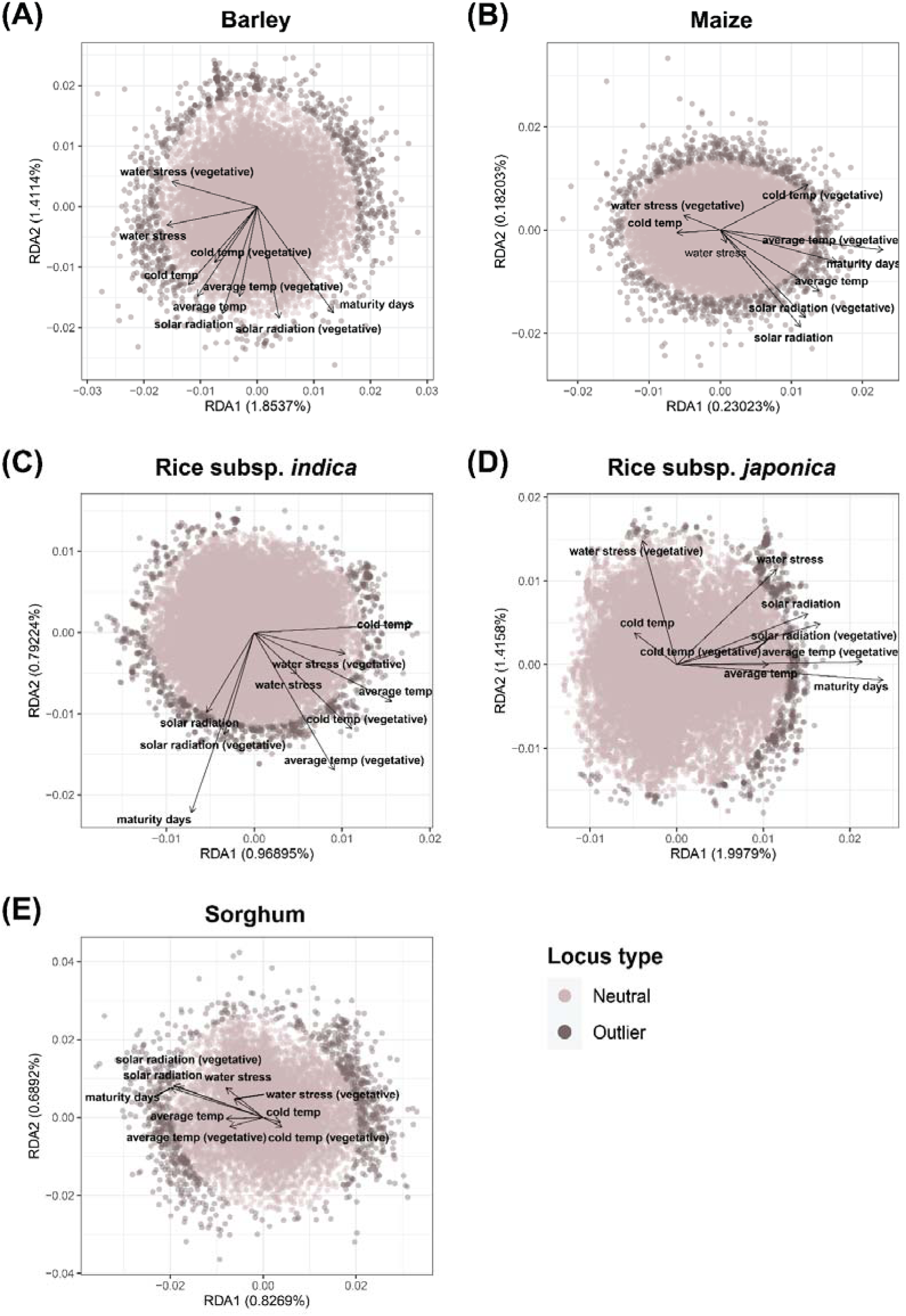
pRDA loading plots for the identification of loci with significant association to *Cycles*-derived environmental variables of the control simulation for use in gradient forest (GF) models. Percent variation explained by the first two pRDA axes (represented as a percentage next to the respective axis) is calculated as the percent variation described by the constrained pRDA axis divided by the variation across all unconstrained axes. Environmentally related (outlier) loci are defined as the 1,000 sites with the most extreme loading along a Mahalanobis distance distribution, calculated between each marker and the center of the first two pRDA axes. For each species, loci identified as “outlier” were retained for use in GF models.

**Figure S3.**
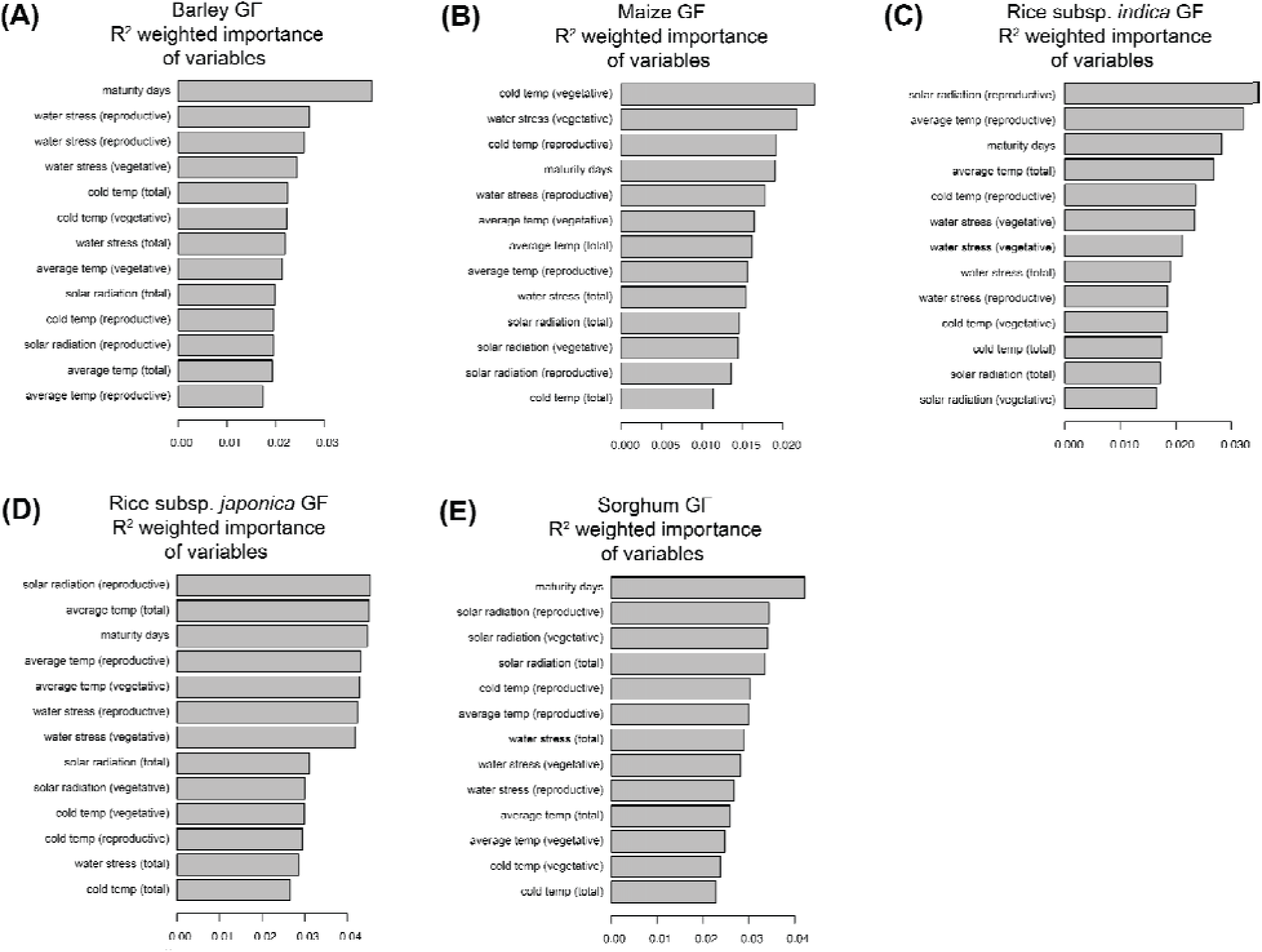
R^2^ importance plots of *Cycles*-derived environmental and growth variables used to build each species’ control GF model for A) barley B) maize C) rice subsp. *indica* D) rice subsp*. japonica* and E) sorghum. Variables are ordered by their relative contribution in describing genome-wide diversity of loci included in each respective GF model.

**Figure S4.**
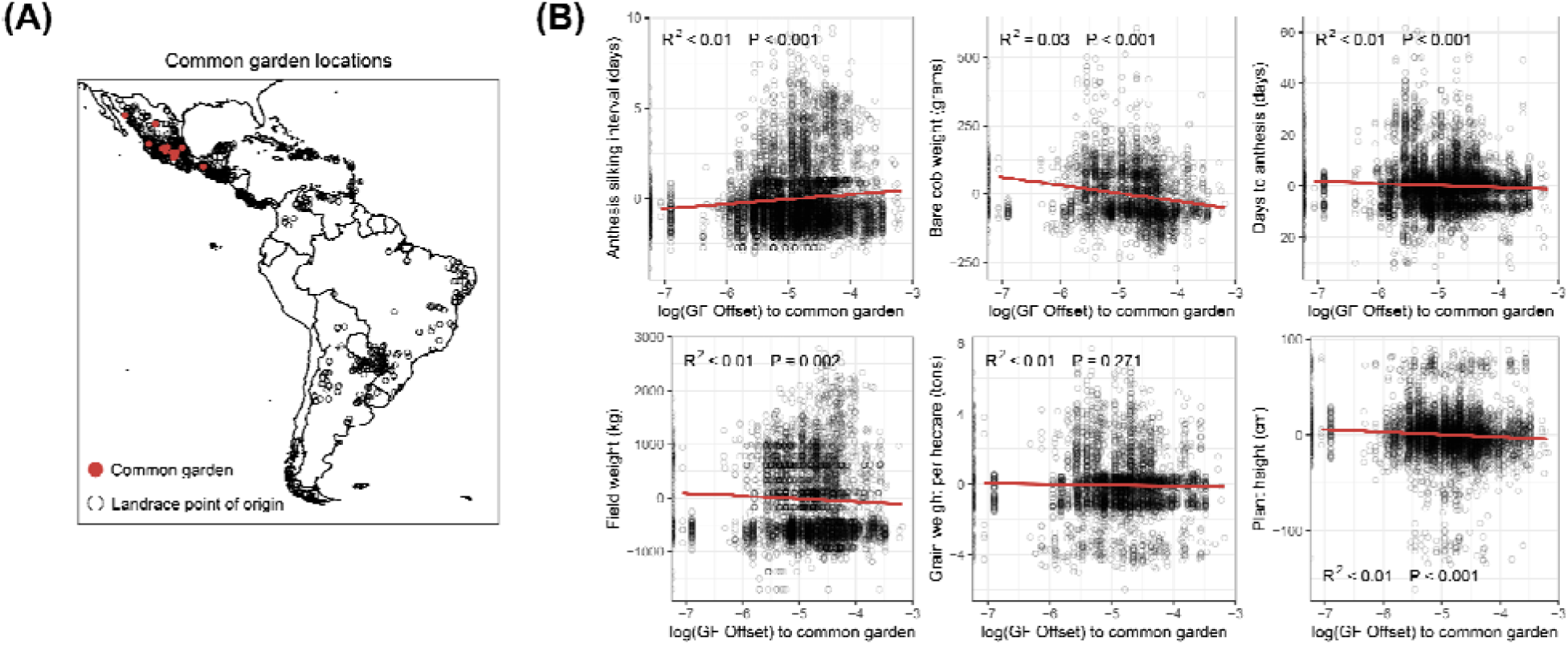
GF models of control genotype-environment associations in maize landraces capture signals of local adaptation. A) Large red points denote sites of common gardens. Black points denote the source locations of landrace accessions grown in common gardens. B) Phenotypic residuals (remaining variation after accounting for experimental design) plotted against the logged GF offset of maize landraces grown in common gardens. GF offset is calculated for each phenotyped accession grown in a common garden as the Euclidean distance of the expected genotype-environment relationship at a common garden common vs the genotype-environment relationship from the landrace accessions’ point of origin. Points with a more negative logged GF offset indicate landrace accessions that are expected to be adapted to conditions at the common garden.

**Figure S5.**
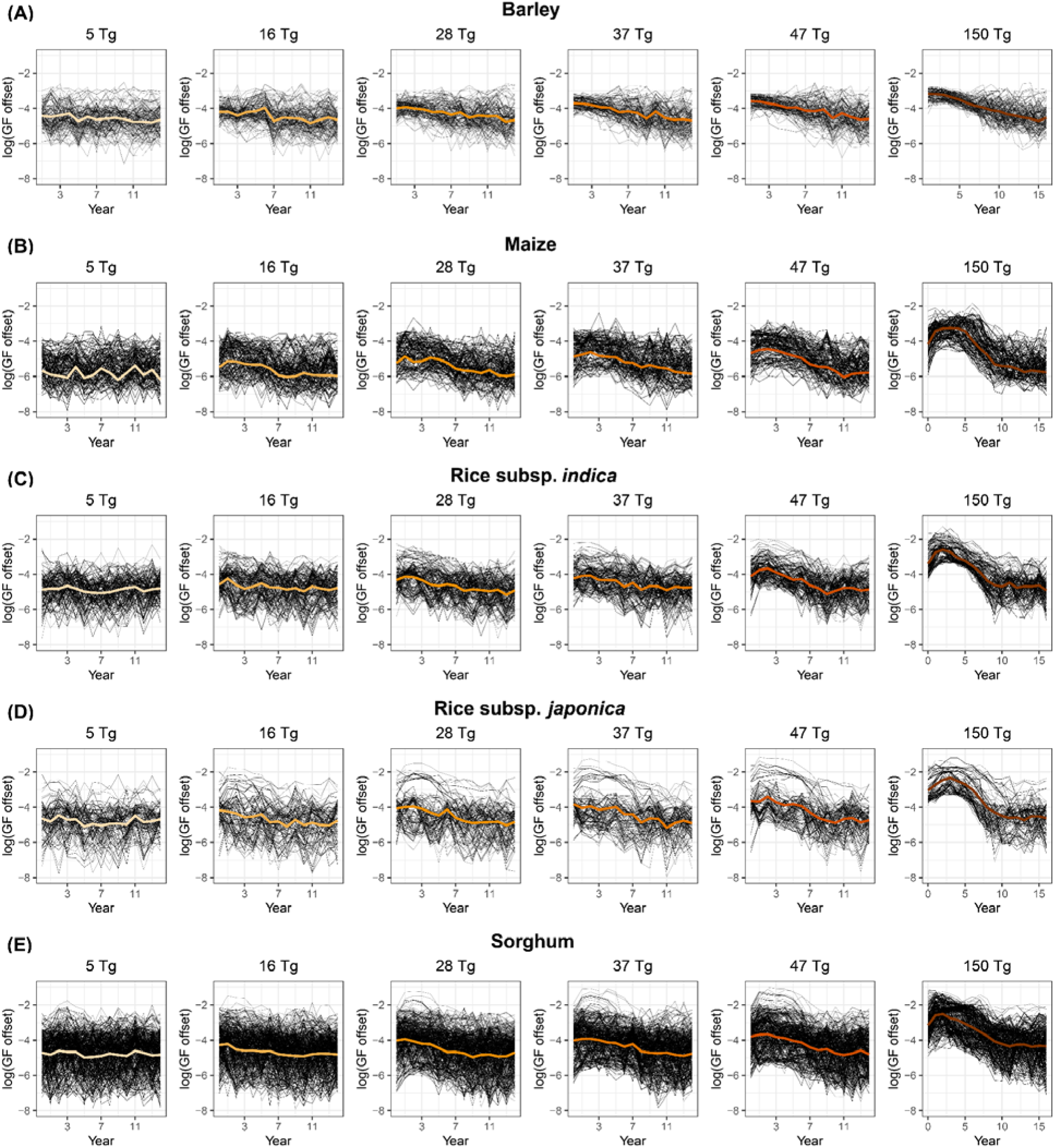
Variation in logged GF offset for A) barley B) maize C) rice subsp. *indica* D) rice subsp*. japonica* and E) sorghum by scenario. Each black line represents the logged GF offset for a modeled landrace accession. The colored line is the average across all individuals by soot scenario. Averaged logged GF offset by soot scenario is the same as shown in Figure 3 B-F.

**Figure S6.**
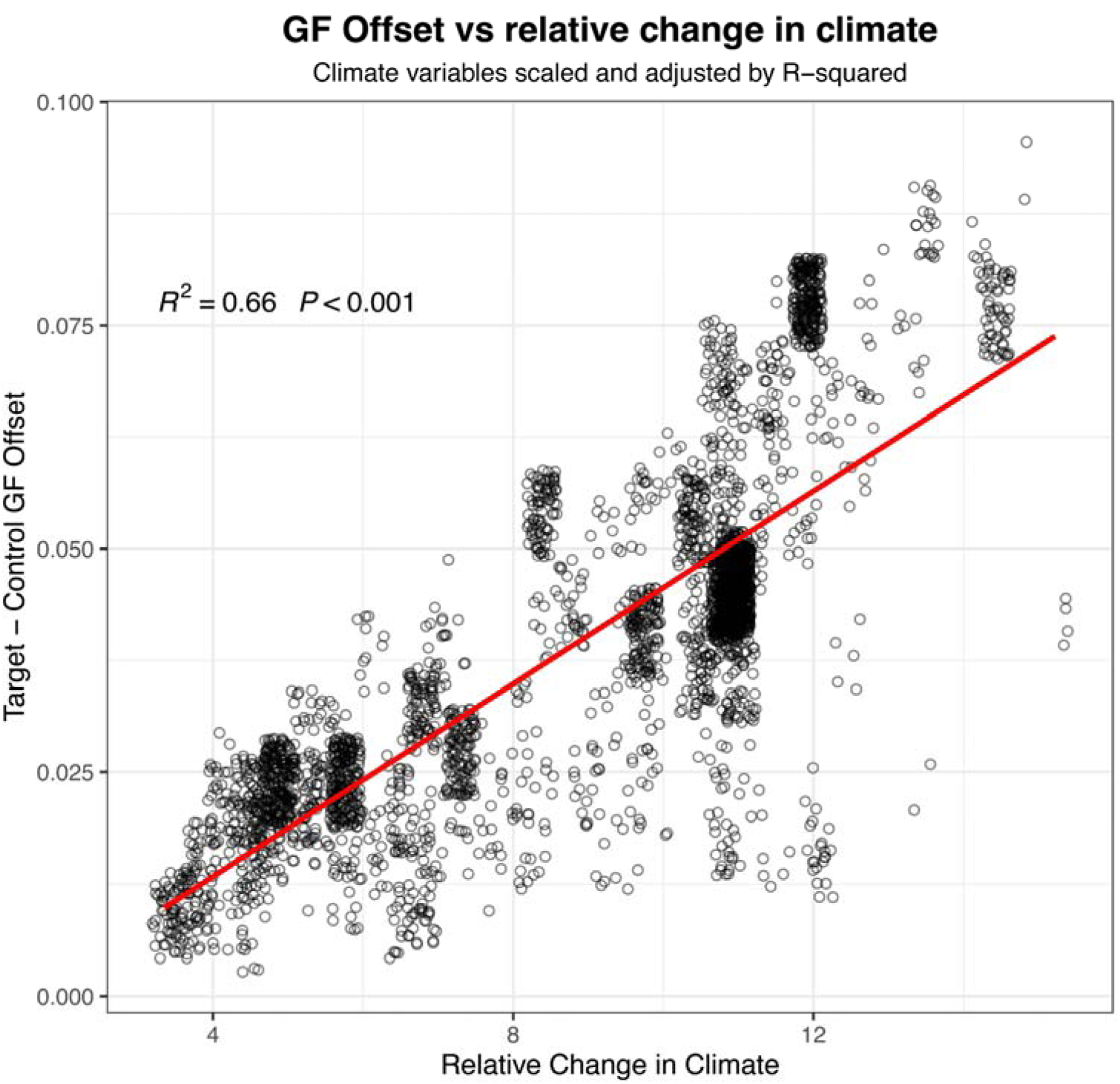
Change in GF offset corresponds to relative changes in climate for maize landrace accessions. Distance between predicted genotype-environment relationships for the 150 Tg “target” scenario and the control scenario vs the environmental change between the 150 Tg target and control scenario.

**Table S1.** Description of environmental and growth variables obtained for *Cycles* simulated landrace accessions (*Cycles*-derived environmental variables). For a given landrace accession, each variable was extracted separately for each simulation year and climate scenario (control, 5 Tg, 6 Tg, 27 Tg, 37 Tg, 47 Tg, 150 Tg).

| Variable | Description |
| --- | --- |
| Average temperature (°C)<br>vegetative, reproductive, cumulative | The daily average temperature for each day crop growth was simulated across the entire growth period (cumulative) or for specific phenological growth stages (vegetative, reproductive). |
| Coldest temperature (°C)<br>vegetative, reproductive, cumulative | The daily minimum temperature for each day crop growth was simulated across the entire growth period (cumulative) or for specific phenological growth stages (vegetative, reproductive). |
| Solar radiation (MJ m<sup>-2</sup>/day)<br>vegetative, reproductive, cumulative | Daily solar radiation for each day crop growth was simulated across the entire growth period (cumulative) or for specific phenological growth stages (vegetative, reproductive). |
| Water Stress (%)<br>vegetative, reproductive, cumulative | Daily water stress for each day crop growth was simulated across the entire growth period (cumulative) or for specific phenological growth stages (vegetative, reproductive). |
| Maturity days | Simulated days to reach physiological maturity. |

**Table S2.** Flowering time genes used in GF models ordered by species. For each flowering time gene, the GeneID for each species’ reference genome is included.

| Species | Gene | GeneID within reference genome |
| --- | --- | --- |
| Maize | *CCA1* | GRMZM2G014902 |
|  | *CCT1* | GRMZM2G381691 |
|  | *CCT11* | GRMZM2G135446 |
|  | *CCT4* | GRMZM2G033962 |
|  | *CCT2* | GRMZM2G004483 |
|  | *CONZ1* | GRMZM2G405368 |
|  | *D8* | GRMZM2G144744 |
|  | *D9* | GRMZM2G024973 |
|  | *DLE1* | GRMZM5G859316 |
|  | *DLF1* | GRMZM2G067921 |
|  | *Gi1* | GRMZM2G107101 |
|  | *Gi2* | GRMZM5G844173 |
|  | *GL15* | GRMZM2G160730 |
|  | *ID1* | GRMZM2G011357 |
|  | *MADS1* | GRMZM2G171365 |
|  | *MADS69* | GRMZM2G171650 |
|  | *PEBP2* | GRMZM2G156079 |
|  | *PEBP24* | GRMZM2G440005 |
|  | *PEBP4* | GRMZM2G075081 |
|  | *PEBP8* | GRMZM2G179264 |
|  | *PRRTF1* | <u>GRMZM2G095727</u> |
|  | *RAP2* | GRMZM2G700665 |
|  | *ZAG6* | GRMZM2G026223 |
|  | *ZCN8* | <u>GRMZM2G019993</u> |
|  | *LHY* | GRMZM2G474769 |
|  | *MADS4* | <u>GRMZM2G032339</u> |
|  | *TOC1a* | GRMZM2G020081 |
| Barley | *CCA1* | Hvcontig_1567295 |
|  | *CEN* | Hvcontig_274284 |
|  | *COL1* | Hvcontig_138334 |
|  | *COL2* | Hvcontig_6805 |
|  | *ELF3* | Hvcontig_80895/67536 |
|  | *EL F4-/ike3* | Hvcontig_42805 |
|  | *FKF1* | Hvcontig_38586 |
|  | *FT* | Hvcontig_54983 |
|  | *GI* | Hvcontig_58270/1580005 |
|  | *GRP7* | Hvcontig_1578172 |
|  | *LHY* | Hvcontig_1567295 |
|  | *LUX* | Hvcontig_2548416 |
|  | *PRR9(5)* | Hvcontig_46739 |
|  | *PRR5(9)* | Hvcontig_41351 |
|  | *PRR7/37* | Hvcontig_94710 |
|  | *TOCI* | Hvcontig_37494 |
|  | *ZTL* | Hvcontig_273830 |
| Rice | *DTH2* | OsR498G0204681300.01 |
|  | *DTH3* | OsR498G0305144600.01 |
|  | *DTH8* | OsR498G0815298200.01 |
|  | *E* | OsR498G1018858200.01 |
|  | *E1* | OsR498G0713935400.01 |
|  | *E2* | OsR498G0713935400.01 |
|  | *E3* | OsR498G0307088600.01 |
|  | *Ehd1* | OsR498G1018858200.01 |
|  | *Ehd2* | OsR498G1018735700.01 |
|  | *Ehd4* | OsR498G0305108900.01 |
|  | *GHd7* | OsR498G0713935400.01 |
|  | *Hd1* | OsR498G0612090700.01 |
|  | *Hd16* | OsR498G0307192700.01 |
|  | *Hd17* | OsR498G0611607800.01 |
|  | *Hd18* | OsR498G0815152000.01 |
|  | *Hd3a* | OsR498G0611656900.01 |
|  | *Se* | OsR498G0612090600.01 |
| Sorghum | *CO* | Sobic.004G007400 |
|  | *CN12* | Sobic.003G295300 |
|  | *CRY1-b1* | Sobic.004G188400 |
|  | *CRY2-2* | Sobic.006G101600 |
|  | *D8* | Sobic.001G120900 |
|  | *Ehd1* | Sobic.010G238700.1 |
|  | *ELF3* | Sobic.009G257300.2 |
|  | *FT1* | Sobic.010G045100 |
|  | *HD6* | Sobic.002G010300 |
|  | *LHY-4* | Sobic.004G279300 |
|  | *Ma2* | Sobic.002G302700 |
|  | *Ma3* | Sobic.001G394400.1 |
|  | *Ma5* | Sobic.001G087100.1 |
|  | *TOC1* | Sobic.004G216700.1 |
|  | *Zfl1* | Sobic.006G201600 |

## Notes

### Competing Interest Statement

The authors have declared no competing interest.

### Summary of Updates

We have edited figures for clarity, and edited text throughout for greater clarity and more accurate interpretation.

## References

1. Lemi, T. Effects of climate change variability on agricultural productivity. Int. J. Environ. Sci. Nat. Resour. 17, (2019).

2. Kemanian, A. R. et al. The Cycles agroecosystem model: Fundamentals, testing, and applications. Comput. Electron. Agric. 227, 109510 (2024).

3. van Klompenburg, T., Kassahun, A. & Catal, C. Crop yield prediction using machine learning: A systematic literature review. Comput. Electron. Agric. 177, 105709 (2020).

4. Altieri, M. A., Nicholls, C. I., Henao, A. & Lana, M. A. Agroecology and the design of climate change-resilient farming systems. Agron. Sustain. Dev. 35, 869–890 (2015).

5. Turco, R. P. et al. Nuclear Winter: Global Consequences Multiple Nuclear Explosions. Science 222, 1283–1292 (1983).

6. Coupe, J., Bardeen, C. G., Robock, A. & Toon, O. B. Nuclear winter responses to nuclear war between the United States and Russia in the whole atmosphere community climate model version 4 and the Goddard institute for space studies ModelE. J. Geophys. Res. 124, 8522–8543 (2019).

7. Jägermeyr, J. et al. A regional nuclear conflict would compromise global food security. Proc. Natl. Acad. Sci. U. S. A. 117, 7071–7081 (2020).

8. Özdoğan, M., Robock, A. & Kucharik, C. J. Impacts of a nuclear war in South Asia on soybean and maize production in the Midwest United States. Clim. Change 116, 373–387 (2013).

9. Xia, L. et al. Global food insecurity and famine from reduced crop, marine fishery and livestock production due to climate disruption from nuclear war soot injection. Nat. Food 3, 586–596 (2022).

10. Lafiandra, D., Riccardi, G. & Shewry, P. R. Improving cereal grain carbohydrates for diet and health. J. Cereal Sci. 59, 312–326 (2014).

11. Azeez, M. A., Adubi, A. O. & Durodola, F. A. Landraces and crop genetic improvement. In Rediscovery of Landraces as a Resource for the Future (InTech, 2018).

12. Villa, T. C. C., Maxted, N., Scholten, M. & Ford-Lloyd, B. Defining and identifying crop landraces. Plant Genet. Resour. 3, 373–384 (2005).

13. Dwivedi, S. L. et al. Landrace germplasm for improving yield and abiotic stress adaptation. Trends Plant Sci. 21, 31–42 (2016).

14. Hoisington, D. et al. Plant genetic resources: what can they contribute toward increased crop productivity? Proc. Natl. Acad. Sci. U. S. A. 96, 5937–5943 (1999).

15. Lasky, J. R., Josephs, E. B. & Morris, G. P. Genotype–environment associations to reveal the molecular basis of environmental adaptation. Plant Cell 35, 125–138 (2023).

16. Vanhove, M. et al. Using gradient Forest to predict climate response and adaptation in Cork oak. J. Evol. Biol. 34, 910–923 (2021).

17. Sklenář, P., Kučerová, J. & Macková, K. Temperature Microclimates Plants Tropical Alpine Environment: How Much does Growth Form Matter? Arctic, Antarctic. Arctic 48, 61–78 (2016).

18. Fitzpatrick, M. C. & Keller, S. R. Ecological genomics meets community-level modelling of biodiversity: mapping the genomic landscape of current and future environmental adaptation. Ecol. Lett. 18, 1–16 (2015).

19. Lasky, J. R. et al. Characterizing genomic variation of Arabidopsis thaliana: the roles of geography and climate. Mol. Ecol. 21, 5512–5529 (2012).

20. Razgour, O. et al. Considering adaptive genetic variation in climate change vulnerability assessment reduces species range loss projections. Proc. Natl. Acad. Sci. U. S. A. 116, 10418–10423 (2019).

21. Savolainen, O., Lascoux, M. & Merilä, J. Ecological genomics of local adaptation. Nat. Rev. Genet. 14, 807–820 (2013).

22. Gutaker, R. M. et al. Genomic history and ecology of the geographic spread of rice. Nat. Plants 6, 492–502 (2020).

23. Lasky, J. R. et al. Genome-environment associations in sorghum landraces predict adaptive traits. Sci. Adv. 1, e1400218 (2015).

24. Rhoné, B. et al. Pearl millet genomic vulnerability to climate change in West Africa highlights the need for regional collaboration. Nat. Commun. 11, 5274 (2020).

25. McLaughlin, C. M. et al. Evidence that variation in root anatomy contributes to local adaptation in Mexican native maize. Evol. Appl. 17, e13673 (2024).

26. Bellis, E. S. et al. Genomics of sorghum local adaptation to a parasitic plant. Proc. Natl. Acad. Sci. U. S. A. 117, 4243–4251 (2020).

27. Gates, D. J., et al. Single-gene resolution of locally adaptive genetic variation in Mexican maize. bioRxiv (2019) doi:10.1101/706739.

28. Rellstab, C., Dauphin, B. & Exposito-Alonso, M. Prospects and limitations of genomic offset in conservation management. Evol. Appl. 14, 1202–1212 (2021).

29. Caproni, L. et al. The genomic and bioclimatic characterization of Ethiopian barley (Hordeum vulgare L.) unveils challenges and opportunities to adapt to a changing climate. Glob. Chang. Biol. 29, 2335–2350 (2023).

30. Láruson, Á. J., Fitzpatrick, M. C., Keller, S. R., Haller, B. C. & Lotterhos, K. E. Seeing the forest for the trees: Assessing genetic offset predictions from gradient forest. Evol. Appl. 15, 403–416 (2022).

31. FAO. Agricultural Production Statistics 2000–2022. https://openknowledge.fao.org/server/api/core/bitstreams/fba4ef43-422c-4d73-886e-3016ff47df52/content (2023) doi:10.4060/cc9205en.

32. Toon, O. B. et al. Rapidly expanding nuclear arsenals in Pakistan and India portend regional and global catastrophe. Sci. Adv. 5, eaay5478 (2019).

33. Shi, Y., Montes, F. & Kemanian, A. R. Cycles[L: A coupled, 3[D, land surface, hydrologic, and agroecosystem landscape model. Water Resour. Res. 59, (2023).

34. Harrison, C. S., et al. A new ocean state after nuclear war. AGU Advances 3, (2022).

35. Russell, J. et al. Exome sequencing of geographically diverse barley landraces and wild relatives gives insights into environmental adaptation. Nat. Genet. 48, 1024–1030 (2016).

36. Romero Navarro, J. A., et al. A study of allelic diversity underlying flowering-time adaptation in maize landraces. Nat. Genet. 49, 476–480 (2017).

37. Hu, Z., Olatoye, M. O., Marla, S. & Morris, G. P. An integrated genotyping-by-sequencing polymorphism map for over 10,000 sorghum genotypes. Plant Genome 12, 180044 (2019).

38. Capblancq, T. & Forester, B. R. Redundancy analysis: A Swiss Army Knife for landscape genomics. Methods Ecol. Evol. 12, 2298–2309 (2021).

39. Breiman, L. Random Forests. Mach. Learn. 45, 5–32 (2001).

40. Ellis, N., Smith, S. J. & Pitcher, C. R. Gradient forests: calculating importance gradients on physical predictors. Ecology 93, 156–168 (2012).

41. Bolaños, J. & Edmeades, G. O. The importance of the anthesis-silking interval in breeding for drought tolerance in tropical maize. Field Crops Res. 48, 65–80 (1996).

42. Bardeen, C. G. et al. Extreme ozone loss following nuclear war results in enhanced surface ultraviolet radiation. J. Geophys. Res. 126, (2021).

43. Kawecki, T. J. & Ebert, D. Conceptual issues in local adaptation. Ecol. Lett. 7, 1225–1241 (2004).

44. Leimu, R. & Fischer, M. A meta-analysis of local adaptation in plants. PLoS One 3, e4010 (2008).

45. Bay, R. A. et al. Genomic signals of selection predict climate-driven population declines in a migratory bird. Science 359, 83–86 (2018).

46. Lachmuth, S., Capblancq, T., Prakash, A., Keller, S. R. & Fitzpatrick, M. C. Novel genomic offset metrics integrate local adaptation into habitat suitability forecasts and inform assisted migration. Ecol. Monogr. 94, (2024).

47. Lasky, J. R., Hooten, M. B. & Adler, P. B. What processes must we understand to forecast regional-scale population dynamics? Proc. Biol. Sci. 287, 20202219 (2020).

48. Glaszmann, J. C. Isozymes and classification of Asian rice varieties. Züchter Genet. Breed. Res. 74, 21–30 (1987).

49. Vigouroux, Y. et al. Selection for earlier flowering crop associated with climatic variations in the Sahel. PLoS One 6, e19563 (2011).

50. Atlin, G. N., Cairns, J. E. & Das, B. Rapid breeding and varietal replacement are critical to adaptation of cropping systems in the developing world to climate change. Glob. Food Sec. 12, 31– 37 (2017).

51. Dahal, K., Li, X.-Q., Tai, H., Creelman, A. & Bizimungu, B. Improving potato stress tolerance and tuber yield under a climate change scenario - A current overview. Front. Plant Sci. 10, 563 (2019).

52. Jain, M., Naeem, S., Orlove, B., Modi, V. & DeFries, R. S. Understanding the causes and consequences of differential decision-making in adaptation research: Adapting to a delayed monsoon onset in Gujarat, India. Glob. Environ. Change 31, 98–109 (2015).

53. Aitken, S. N. & Whitlock, M. C. Assisted gene flow to facilitate local adaptation to climate change. Annu. Rev. Ecol. Evol. Syst. 44, 367–388 (2013).

54. Marsh, D. R. et al. Climate change from 1850 to 2005 simulated in CESM1(WACCM). J. Clim. 26, 7372–7391 (2013).

55. Toon, O. B., Turco, R. P., Westphal, D., Malone, R. & Liu, M. A multidimensional model for aerosols: Description of computational analogs. J. Atmos. Sci. 45, 2123–2144 (1988).

56. Bardeen, C. G., Toon, O. B., Jensen, E. J., Marsh, D. R. & Harvey, V. L. Numerical simulations of the three-dimensional distribution of meteoric dust in the mesosphere and upper stratosphere. J. Geophys. Res. (2008) doi:10.1029/2007JD009515.

57. Bardeen, C. G., Garcia, R. R., Toon, O. B. & Conley, A. J. On transient climate change at the Cretaceous-Paleogene boundary due to atmospheric soot injections. Proc. Natl. Acad. Sci. U. S. A. 114, E7415–E7424 (2017).

58. Robock, A., Oman, L. & and Stenchikov, G. L. Nuclear winter revisited with a modern climate model and current nuclear arsenals: Still catastrophic consequences. J. Geophys. Res. 112, (2007).

59. Hengl, T. et al. Mapping soil properties of Africa at 250 m resolution: Random forests significantly improve current predictions. PLoS One 10, e0125814 (2015).

60. Leonard, L. & Duffy, C. J. Automating data-model workflows at a level 12 HUC scale: Watershed modeling in a distributed computing environment. Environ. Model. Softw. 61, 174–190 (2014).

61. Leonard, L. & Duffy, C. J. Essential Terrestrial Variable data workflows for distributed water resources modeling. Environ. Model. Softw. 50, 85–96 (2013).

62. Leonard, L. & Duffy, C. Visualization workflows for level-12 HUC scales: Towards an expert system for watershed analysis in a distributed computing environment. Environ. Model. Softw. 78, 163–178 (2016).

63. Capblancq, T., Luu, K., Blum, M. G. B. & Bazin, E. Evaluation of redundancy analysis to identify signatures of local adaptation. Mol. Ecol. Resour. 18, 1223–1233 (2018).

64. Oksanen, J. et al. vegan community ecology package version 2.6-2 April 2022. The Comprehensive R Archive Network. Available online: http://cran.r-project.org *(accessed on* 15 August 2022) (2022).

65. Huang, M., Liu, X., Zhou, Y., Summers, R. M. & Zhang, Z. BLINK: a package for the next level of genome-wide association studies with both individuals and markers in the millions. Gigascience 8, (2019).

66. Ross-Ibarra, J. GTF and GFF for maize. figshare 10.6084/M9.FIGSHARE.895628.V1 (2014).

67. Du, H. et al. Sequencing and de novo assembly of a near complete indica rice genome. Nat. Commun. 8, 15324 (2017).

68. McCormick, R. F. et al. The Sorghum bicolor reference genome: improved assembly, gene annotations, a transcriptome atlas, and signatures of genome organization. Plant J. 93, 338–354 (2018).

69. Purcell, S. et al. PLINK: a tool set for whole-genome association and population-based linkage analyses. Am. J. Hum. Genet. 81, 559–575 (2007).

70. Hahsler, M., Piekenbrock, M. & Doran, D. dbscan: Fast Density-Based Clustering with R. J. Stat. Softw. 91, (2019).

71. Hijmans, R.J., Williams, E., and Vennes, C. geosphere: Spherical Trigonometry. R package version 1.5-14. https://CRAN.R-project.org/package=geosphere (2022).

